# Accelerating antibiotic discovery from Earth’s isolated habitats through protein language models and microfluidics

**DOI:** 10.1101/2025.11.13.688364

**Authors:** Zixin Kang, Qize Zhou, Haohong Zhang, Jiayi Liu, Kun Liu, Kouyi Zhou, Yufei Zhang, Yuting Wang, Peng Chen, Bi-Feng Liu, Kang Ning

## Abstract

The antibiotic potential of Earth’s isolated habitat microbiomes remains largely unexplored. Accessing this resource requires reliable prioritization across divergent peptide sequences and efficient experimental validation. Here we present MicroSPEAR, an end-to-end framework for antimicrobial peptide (AMP) discovery integrating SPEAR, a sequence-structure-aware ensemble predictor, with MicroGrad, a microfluidic testing platform. SPEAR generalized to low-homology peptides and prioritized potent candidates despite training on binary activity labels. MicroGrad simplified liquid handling and shortened susceptibility testing from approximately 18 h to 4 h. Applied to a catalogue derived from 4,480 metagenomes spanning 15 isolated habitat types, SPEAR identified 383,027 non-redundant candidate AMPs, 63.1% of which lacked detectable homologues in reference databases. Of 85 synthesized candidates, 79 exhibited antibacterial activity at ≤64 μmol l□^1^. Mechanistic analyses revealed distinct modes of membrane disruption. Four broad-spectrum candidates with low toxicity significantly reduced bacterial burdens in a mouse thigh infection model. MicroSPEAR connects underexplored microbial diversity with experimentally validated antibiotic leads and provides a foundation for lab-in-the-loop discovery.

## Introduction

Antimicrobial resistance (AMR) threatens the effective treatment of bacterial infections and creates an urgent need for new antibiotic agents [1–3]. An estimated 1.14 million deaths worldwide were directly attributable to bacterial AMR in 2021, with annual deaths projected to reach 1.91 million by 2050 under the reference scenario [3]. Antimicrobial peptides (AMPs) represent a promising source of antibiotic agents owing to their diverse mechanisms of antibacterial action and potentially lower propensity to induce resistance [4]. Advances in metagenomic sequencing and assembly have enabled the discovery of candidate AMPs encoded by small open reading frames (smORFs) directly from environmental samples [5, 6]. Coupled with computational prediction, metagenomic mining has identified millions of candidate AMPs and yielded hundreds of experimentally validated peptides [7–9]. Yet extending systematic antibiotic discovery into broader, uncharacterized sequence spaces confronts two fundamental challenges: sequence novelty and experimental validation scale.

On the computational side, machine learning has become central to the identification and prioritization of AMPs from these large sequence collections [4, 10, 11]. Classifier models such as Macrel [12] and amPEPpy [13] use engineered sequence descriptors, whereas deep learning models such as AMP Scanner v2 [14] and AMPlify [15] learn representations directly from primary sequences. Nevertheless, screening poorly characterized microbiomes requires reliable predictions for sequences that may differ substantially from those represented in training data. Under such distribution shifts, maintaining high sensitivity while limiting false positives remains essential, particularly when the number of predicted candidates greatly exceeds experimental capacity. Effective prioritization must also distinguish highly potent peptides from weakly active ones, motivating frameworks that couple out-of-distribution AMP recognition with potency-aware ranking.

On the experimental side, high-throughput validation is equally critical for translating computational predictions into verified leads [10]. Conventional broth microdilution assays require serial dilution, bacterial inoculation and typically overnight incubation to determine minimum inhibitory concentrations (MICs) [16–18], constraining the rate at which candidate peptides can be evaluated. Although microfluidic platforms have accelerated antimicrobial susceptibility testing (AST) for established small-molecule antibiotics by miniaturizing assays and simplifying sample processing [16, 19, 20], their systematic integration into large-scale AMP discovery workflows remains limited. Overcoming the dual challenges of sequence novelty and experimental scale therefore calls for an integrated computational-experimental framework that couples low-homology recognition and potency-aware ranking with rapid, automated susceptibility testing.

Deploying such a framework to uncover genuinely divergent antibiotic scaffolds requires looking beyond well-sampled environments [21]. Historical discoveries (such as paclitaxel [22] from the Pacific yew, cyclosporin A [23] from a Norwegian soil fungus, rapamycin [24] from an Easter Island soil bacterium and ziconotide [25] from cone-snail venom) illustrate how biogeographically distinct niches yield clinically valuable molecules. Here, we operationally defined isolated habitats as environments where geographic separation, physical barriers or pronounced physicochemical filtering constrain microbial dispersal and establishment [26–32] (Supplementary Table 1, 2). Such biogeographic partitioning drives evolutionary divergence across microbial communities and their gene repertoires, creating an underexplored reservoir for discovering novel antibiotics.

Recent global sampling campaigns and genome-resolved surveys have now made these remote ecosystems accessible to large-scale metagenomic mining. For example, the Vanishing Glaciers Project [33] has surveyed glacier-fed streams across the world’s major mountain ranges, while the MOSAiC expedition [34] and the Antarctic Circumnavigation Expedition (ACE) [35] have generated metagenomic resources from Arctic and Southern Ocean ecosystems. The Census of Deep Life (CoDL) [36] and the Mariana Trench Environment and Ecology Research (MEER) project [37] have further supported investigations of subsurface and hadal microbial communities. Together, these initiatives offer the opportunity to move from historical serendipitous discovery to systematic bioprospecting across Earth’s isolated habitats.

In this study, we present MicroSPEAR, an end-to-end framework that couples SPEAR, a sequence-structure-aware ensemble predictor, with MicroGrad, a microfluidic AST platform. In homology-controlled evaluations, SPEAR retains specificity at high recall as sequence identity between training and test sets decreases, and corrects a substantial fraction of errors shared by established predictors. On an AMP-like peptide test set compiled from multiple studies, it outperforms the current state-of-the-art (SOTA) baseline, improving the Matthews correlation coefficient (MCC) by 6.8% and reducing the Brier score by 19.0% on AMP prediction. Its rankings also show 10.6% stronger Spearman correlation with antimicrobial potency than the baseline. MicroGrad addresses the experimental component by automating concentration-gradient generation and enabling direct visual readout, reducing susceptibility-testing time from approximately 18 h with conventional broth microdilution to about 4 h.

We further applied SPEAR to the Global Isolated-Habitat Microbiome smORF Catalog (GIHMsC), which we constructed from 4,480 metagenomes spanning 15 isolated habitat types. This analysis identified 383,027 non-redundant candidate AMPs encoded by microorganisms spanning 159 bacterial and 23 archaeal phyla, with 63.1% lacking detectable homologues in reference databases. Of 85 synthesized candidates evaluated across an eight-strain pathogen panel, 79 exhibited antibacterial activity against at least one tested strain at concentrations ≤64 μmol l□^1^. Among the 32 top-ranked candidates (SPEAR score > 0.95) tested, 15 exhibited high potency, with MICs ≤16 μmol l□^1^ against at least four strains or ≤8 μmol l□^1^ against at least two strains. Furthermore, four broad-spectrum candidates with low toxicity reduced bacterial burdens by two to four orders of magnitude in a mouse thigh infection model, supporting their potential as anti-infective leads. By integrating robust computational prioritization with rapid experimental validation, MicroSPEAR provides a foundation for lab-in-the-loop antibiotic discovery.

## Results

### GIHMsC reveals underexplored, habitat-partitioned peptide space

We first sought to establish a global genome-resolved data resource to explore the diversity and distribution of microbial peptides across Earth’s isolated habitats. To this end, we compiled 153,243 metagenome-assembled genomes (MAGs) from 4,480 metagenomes across 2,428 sampling locations spanning 15 habitat types (Fig. 1a, b). These habitats were characterized using a standardized framework of environmental isolation and associated physicochemical features, including temperature, salinity, water availability and oxygen availability (Supplementary Tables 1 and 2). The dataset spans a combined elevational and depth range of approximately 15,000 m and a temperature range of approximately 120 °C, including more than 400 high-altitude freshwater, saline lake, glacier and hot spring samples from the Qinghai-Tibet Plateau [6, 38]. Following genome-quality assessment with CheckM2 [39], we retained 119,107 MAGs meeting our completeness (≥50%) and contamination (≤5%) thresholds (Supplementary Fig. 1).

**Figure 1.**
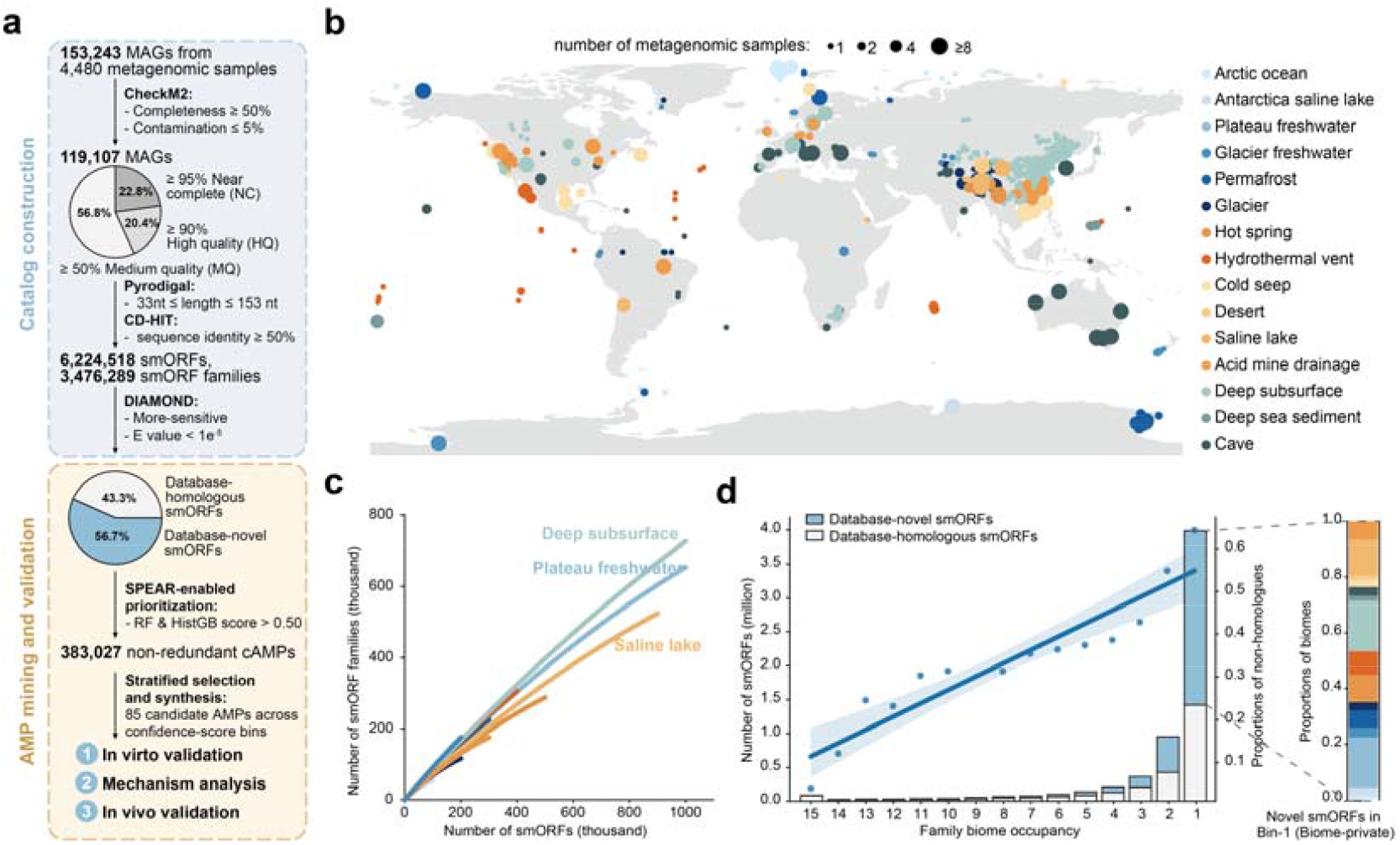
Global cataloguing and SPEAR-guided discovery of antimicrobial peptides (AMPs) from isolated-habitat microbiomes. **a**, End-to-end workflow for AMP bioprospecting. A total of 153,243 metagenome-assembled genomes (MAGs) were recovered from 4,480 metagenomic samples, of which 119,107 passed CheckM2 quality filtering. Pyrodigal predicted 6,224,518 small open reading frames (smORFs), which were grouped into 3,476,289 sequence families using CD-HIT. DIAMOND searches against reference protein databases identified 3,527,632 database-novel smORFs with no detectable homologues. SPEAR-based prioritization yielded 383,027 non-redundant candidate AMPs (cAMPs), from which 85 were selected across prediction-score bins for synthesis and experimental validation. Downstream experiments included in vitro validation, mechanistic characterization and in vivo validation. Pie charts show the quality distribution of retained MAGs and the proportions of database-homologous and database-novel smORFs. **b**, Geographic distribution of metagenomic samples. Dot size indicates the number of samples at each geographic location. **c**, smORF family accumulation curves across isolated habitats. The number of smORF families continued to increase with the number of sampled smORFs, with no clear plateau observed, indicating that smORF diversity in isolated-habitat microbiomes remains far from saturation. **d**, Habitat breadth of smORF families. Most non-homologous smORF families were detected in only one habitat type. The stacked bar plot shows the habitat-source composition of non-homologous and habitat-specific smORFs.

From this curated genome collection, we constructed the Global Isolated-Habitat Microbiome smORF Catalog (GIHMsC), comprising 6,224,518 smORFs predicted to encode peptides of 10-50 amino acids. Plateau freshwater contributed the largest number of smORFs (1,246,732), followed by deep subsurface environments (1,063,322) and saline lakes (947,178). Clustering at 50% amino acid identity and 95% alignment coverage resolved 3,476,289 operational families, revealing extensive sequence diversity even at this permissive identity threshold. Searches against reference protein and smORF databases identified 3,527,632 smORFs (56.7%) without detectable homologs under our search criteria, hereafter termed database-novel smORFs (Methods; Fig. 1a). Furthermore, 99.41% of these database-novel smORFs lacked detectable sequence similarity to known reference AMPs (Supplementary Fig. 2a), indicating that more than half of the catalogue represents predicted small-protein sequences absent from existing protein and antimicrobial peptide resources. Despite this extensive diversity, family accumulation curves continued to rise with increasing numbers of sampled smORFs, without approaching an asymptote (Fig. 1c). These results indicate that the sampled sequence space remains far from saturated and support deeper exploration of isolated-habitat microbiomes to recover additional smORF families.

Beyond this unsaturated diversity within habitats, smORF repertoires were strongly partitioned among habitats. More than 4 million smORFs belonged to families detected in only one habitat type within GIHMsC, hereafter termed biome-private families. The fraction of database-novel smORFs increased as family habitat breadth decreased, indicating that narrowly distributed families contributed disproportionately to the uncataloged sequence space (Fig. 1d). At the opposite extreme, only 0.014% of smORFs belonged to families detected across all 15 habitat types, and 96.2% of these broadly distributed smORFs had homologs among known proteins. This contrast reveals a small, broadly shared and largely database-represented core alongside a much larger, habitat-partitioned reservoir enriched in previously unrepresented sequences. Together with the unsaturated family accumulation curves, these findings support both deeper sampling within isolated habitats and broader sampling across habitat types to expand peptide discovery.

### SPEAR corrects consensus errors and improves generalization

To mine the novel sequence space of GIHMsC, we developed SPEAR, a protein language model (pLM)-based framework for AMP prediction. We trained and evaluated SPEAR on a non-redundant dataset of 13,652 peptides (6,169 AMPs and 7,483 non-AMPs; 10-50 amino acids) compiled from DBAASP [40], DRAMP [41], UniProt [42], a published benchmark [43] and six experimental studies (Methods). Contact-map clustering of ESMFold-predicted structures across the 6,169 positive AMPs resolved nine structural archetypes spanning helical, mixed α/β, β-rich and extended conformations that were not uniquely specified by individual physicochemical descriptors (Supplementary Figs. 3 and 4 and Supplementary Table 3). Benchmarking peptide representations across the dataset further showed that ESM2 [44] sequence embeddings best preserved local AMP/non-AMP label homogeneity, whereas structure-aware SaProt [45] embeddings best separated the nine structural archetypes (Supplementary Fig. 5 and Supplementary Tables 4 and 5). To capture both complementary views, SPEAR concatenates ESM2 and SaProt embeddings and feeds the fused representation into separate random forest and histogram-based gradient boosting classifiers, whose predicted probabilities are averaged to produce the final score (Fig. 2a). To benchmark SPEAR against existing approaches, we retrained and evaluated five representative AMP predictors, including an engineered descriptor-based model (amPEPpy), task-specific sequence representation learning models (AMP Scanner v2 and EvoGradient), and pLM-based models (HMD-AMP and AMP-BERT).

**Figure 2.**
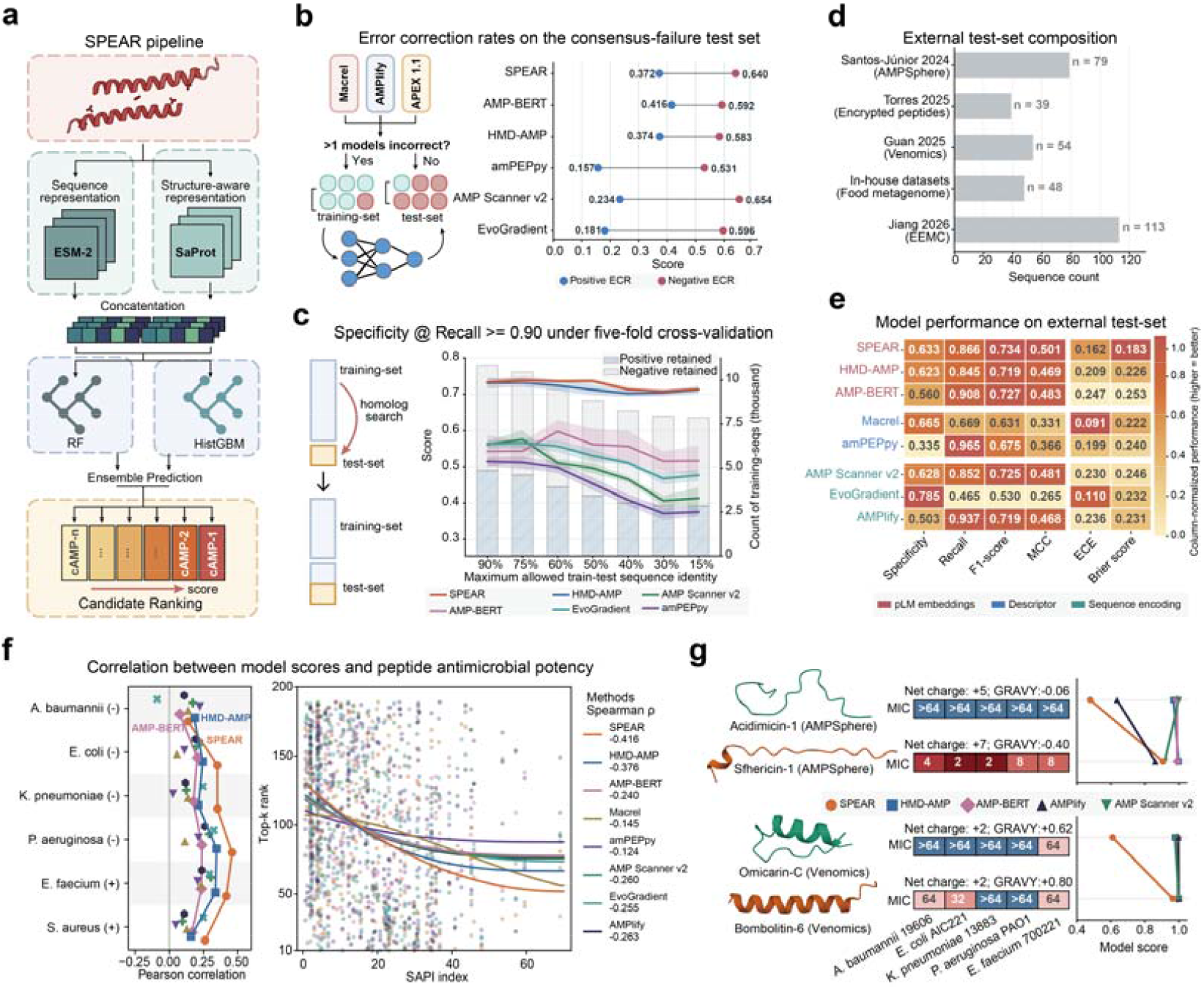
Sequence-structure-informed SPEAR enables robust AMP prediction across divergent sequence space. **a**, Overview of the SPEAR framework. **b**, Error-correction performance on the consensus-failure test set. The test set comprised peptides misclassified by a majority vote of Macrel [12], AMPlify [15] and APEX 1.1 □46]. Overall, error-correction rates (ECRs) quantify the fractions of these errors corrected by each evaluated model across the test set. **c**, Specificity at recall□≥□0.90 under identity-controlled five-fold cross-validation. Within each fold, the test set was held fixed, and training sequences whose maximum identity to any test sequence was at or above the indicated cutoff were excluded. Bars show the numbers of AMP and non-AMP training sequences retained at each train-test sequence-identity cutoff. **d-g**, Benchmarking SPEAR against seven baseline models (HMD-AMP, AMP-BERT, Macrel, amPEPpy, AMP Scanner v2, EvoGradient, and AMPlify) on an independent external test set comprising 333 experimentally characterized peptides. Peptides were classified as AMP, weak-AMP, or non-AMP according to their experimentally measured MIC profiles. **d**, Composition of the external test set. With the exception of Macrel, which was used in the construction of the AMPSphere database, none of the benchmarked models contributed to test-set construction. **e**, External-test performance of SPEAR and baselines. Weak-AMPs and non-AMPs were jointly treated as the negative class. Models are grouped by peptide representation: protein language model (pLM) embeddings, engineered sequence descriptors, or sequence representations learned from scratch. Performance was assessed using specificity, recall, F1-score, Matthews correlation coefficient (MCC), expected calibration error (ECE), and Brier score. Lower ECE and Brier scores indicate better calibrated and more accurate probabilistic predictions, respectively. For visual comparability, values were normalized within each metric so that darker shading denotes better performance. **f**, Association between model predictions and peptide antimicrobial potency. Left, Pearson correlations between model prediction scores and MIC-derived antimicrobial potency against six bacterial species. Sample sizes (−) and (+) denote Gram-negative and Gram-positive bacteria, respectively. Right, relationship between peptide ranks based on descending model scores (rank 1 indicates the highest score) and the spectrum-adjusted potency index (SAPI). To assess the ability of models to prioritize more potent AMPs, non-AMP peptides were excluded from the analysis. Each point represents a peptide ranked by a given model, curves show fitted trends, and Spearman correlation coefficients (ρ) are provided in the legend. **g**, Representative matched-pair cases illustrating SPEAR’s prioritization of highly potent AMPs. ESMFold-predicted structures are shown on the left. Heatmaps in the center show MICs (μmol l□^1^) against four Gram-negative strains (*A. baumannii* ATCC 19606, *E. coli* ATCC 25922, *K. pneumoniae* ATCC 13883, and *P. aeruginosa* PAO1) and one Gram-positive strain (*E. faecium* ATCC 700221), with values >64 μmol l□^1^ indicating no detectable activity within the tested range. The right panels compare AMP scores assigned by SPEAR and five baseline models.

We first asked whether SPEAR could correct prediction errors made by multiple established methods (Fig. 2b). In large-scale peptide mining, such shared false positives can inflate candidate lists and increase the burden of downstream peptide synthesis and activity testing. Using three widely used predictors, Macrel [12], AMPlify [15] and APEX 1.1 [46] as fixed references, we constructed a baseline consensus-failure (BCF) test set comprising peptides misclassified by at least two references, yielding 2,871 AMPs and 228 non-AMPs (Methods). SPEAR and five representative models were re-trained on the remaining data and evaluated on the BCF test set using the error-correction rate (ECR). We focused particularly on negative ECR, which quantifies the elimination of shared false positives and reflects the potential to reduce fruitless synthesis. SPEAR corrected 64.0% of consensus false positives, exceeding HMD-AMP and AMP-BERT by 5.7 and 4.8 percentage points, respectively (Fig. 2b). SPEAR also recovered 37.2% of consensus false negatives, comparable to HMD-AMP (37.4%). Compared with AMP Scanner v2, SPEAR achieved a similar false-positive correction rate (64.0% versus 65.4%) while recovering more missed AMPs (37.2% versus 23.4%). These results demonstrate that SPEAR suppresses consensus false positives while recovering bona fide AMPs missed by established tools.

Next, we evaluated model specificity at high recall in sequence-out-of-distribution settings. We performed five-fold cross-validation with fixed test folds and progressively stricter train-test sequence-identity thresholds (90%-15%). For each threshold, training peptides exceeding the permitted identity with any test peptide were removed (Fig. 2c). Performance was evaluated using specificity at recall ≥0.90 (Spec@R90), which measures the ability to reject non-AMPs while retaining high sensitivity to AMPs. As the maximum permitted identity decreased, SPEAR maintained the highest Spec@R90, declining only from 0.733 to 0.708, compared with 0.731 to 0.704 for the SOTA baseline, HMD-AMP. In contrast, Spec@R90 decreased more substantially for amPEPpy, from 0.515 to 0.371, and AMP Scanner v2, from 0.561 to 0.406. EvoGradient declined from 0.563 to 0.477, whereas AMP-BERT fluctuated between 0.516 and 0.597. These results highlight SPEAR’s robustness and generalization to peptides with low sequence similarity to the training data.

Additionally, we performed ablation experiments across 150 representation-classifier configurations under the same five-fold cross-validation setup with a 30% train-test sequence-identity cutoff (Supplementary Fig. 6). With the representation module held fixed, replacing SPEAR’s tree-based ensemble with an individual random-forest classifier decreased Spec@R90 from 0.709 to 0.645, and to 0.661 and 0.560 for the multilayer perceptron and logistic regression classifiers, respectively. With the classifier module held fixed, using sequence-only or structure-aware-only pLM representations reduced the average Spec@R90 to 0.694 and 0.649, respectively. Overall, SPEAR achieved a Spec@R90 improvement of 0.111 over the protein function prediction baseline (logistic regression based on ESM2-650M representations). These findings indicate that both the fusion of sequence and structure-aware representations and tree-classifier ensembling contribute to SPEAR’s generalization to low-homology peptides.

### SPEAR enables potency-informed AMP prioritization

To evaluate SPEAR’s performance on prioritization of highly potent candidates, we assembled an external test set comprising 333 AMP-like peptides derived from human and other animal proteomes and microbial genomes. These peptides were collected from four published studies [9, 47–49] and an in-house dataset (Fig. 2d, Methods). This dataset encompasses 1,980 experimentally measured MICs across six bacterial species, categorizing the peptides into non-AMPs (n = 144), weak-AMPs (n = 47) and active AMPs (n = 142; Methods).

To evaluate the ability to prioritize peptides active against multiple pathogens, we treated AMPs as positives and grouped weak-AMPs with non-AMPs as negatives. SPEAR achieved the highest F1 score (0.734) and MCC (0.501), together with the lowest Brier score (0.183) among the eight evaluated predictors (Fig. 2e). SPEAR combined a recall of 0.866 with a specificity of 0.633. By comparison, amPEPpy and AMPlify achieved higher recall (0.965 and 0.937) but lower specificity (0.335 and 0.503), whereas EvoGradient achieved the highest specificity (0.785) at a recall of only 0.465. Relative to HMD-AMP, SPEAR improved MCC by 6.8% and lowered the Brier score by 19.0%, demonstrating superior binary discrimination and probability calibration.

We further defined a spectrum-adjusted potency index (SAPI) to quantify antimicrobial potency across tested bacterial species (Methods). To assess potency-based prioritization, we excluded non-AMPs and ranked the remaining 189 peptides by each model’s prediction scores, with rank 1 assigned to the highest-scoring candidate. We then assessed the association between ranking position and SAPI using Spearman correlation. SPEAR showed the strongest inverse association (ρ = −0.416), followed by HMD-AMP (ρ = −0.376). Associations were weaker for the remaining baselines, with ρ ranging from −0.263 to −0.124 (Fig. 2f). To further illustrate differences in potency-based prioritization, we examined two pairs of length-matched peptides with similar physicochemical properties but contrasting antimicrobial activities from the full external test set (Fig. 2g). In the AMPSphere pair [9], Acidimicin-1 had MIC values >64 μmol l□^1^ against all five tested strains, whereas Sfhericin-1 had MIC values of 2-8 μmol l□^1^. In the Venomics pair [48], Bombolitin-6 showed greater activity against E. coli (MIC, 32 versus >64 μmol l□^1^) and A. baumannii (64 versus >64 μmol l□^1^). Across both pairs, SPEAR assigned substantially higher scores to the more active peptide: 0.906 versus 0.479 for Sfhericin-1 and Acidimicin-1, and 0.999 versus 0.611 for Bombolitin-6 and Omicarin-C. By comparison, baseline predictors assigned near-maximal scores (~1.0) to both active and inactive peptides in each pair. Together with the aggregate ranking analysis, these results support the use of SPEAR scores to prioritize peptides with greater antimicrobial potency.

To evaluate species-level potency prioritization, we calculated Pearson correlations between model prediction scores and −log□(MIC) across individual pathogens. SPEAR showed positive correlations across all six species and achieved the highest correlation coefficients for five of them: *P. aeruginosa* (*r* = 0.460), *E. faecium* (*r* = 0.418), *E. coli* (*r* = 0.352), *K. pneumoniae* (*r* = 0.352), and *S. aureus* (*r* = 0.261), alongside *r* = 0.139 for *A. baumannii* (Fig. 2f). These results demonstrate that SPEAR scores capture antimicrobial potency across diverse Gram-negative and Gram-positive pathogens.

### Diverse microbial lineages encode candidate AMPs

To characterize the AMP-encoding potential of Earth’s isolated habitat microbiomes, we screened 4,738,802 smORFs from GIHMsC with SPEAR after collapsing redundant sequences (Methods). Whereas existing predictors showed limited candidate overlap and pronounced cross-model disagreement across GIHMsC families of varying habitat breadth (Supplementary Fig. 2b, c), SPEAR identified 383,027 candidate AMPs (cAMPs), 63.1% of which were encoded by database-novel smORFs (Fig. 3a). Of these, 2,513 had a mean SPEAR score >0.95, representing 0.053% of the screened smORFs. Candidates were distributed across diverse bacterial and archaeal lineages, as illustrated by a family-level phylogeny constructed using the GTDB-Tk backbone (Fig. 3b). Pseudomonadota and Bacteroidota contributed the largest numbers of candidates, with 74,827 and 66,555 peptides encoded by members of 501 and 264 taxonomic families, respectively. Substantial contributions also came from Actinomycetota (24,297), Desulfobacterota (17,552) and Chloroflexota (16,794). The candidate repertoire thus extended across numerous lineages within this largely uncultivated bacterial radiation.

**Figure 3.**
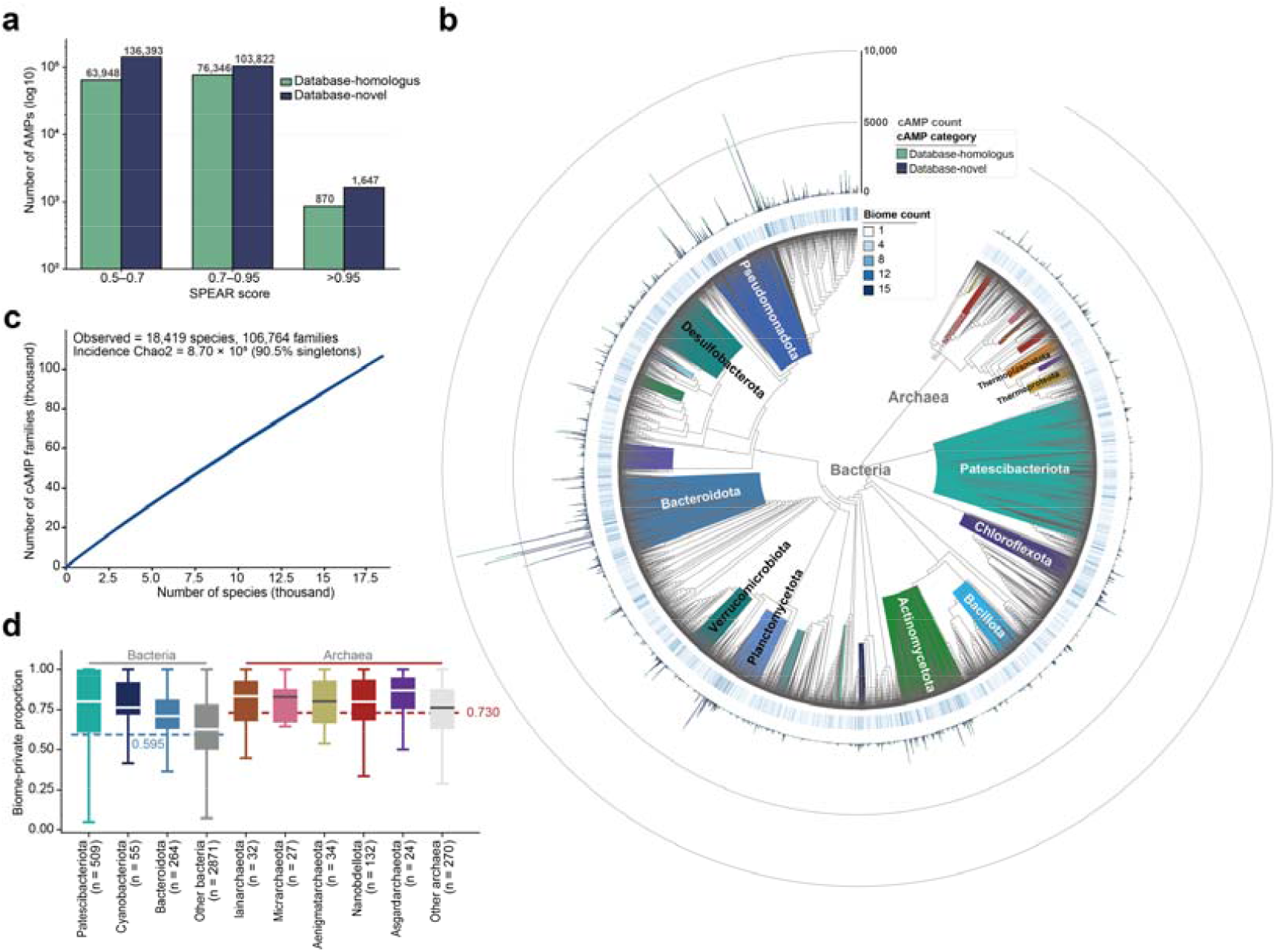
Taxonomic distribution, unsaturated diversity and habitat partitioning of SPEAR-prioritized candidate AMPs. **a**, Counts of non-redundant candidate AMPs (n = 383,027) in three mean SPEAR-score intervals (0.50 to 0.70, 0.70 to 0.95 and >0.95), shown on a log10 scale and partitioned into database-homologous and database-novel smORFs. Candidates required scores above 0.5 from both SPEAR classifiers. **b**, Family-level phylogeny of candidate-encoding taxa placed on the GTDB reference backbone. Inner sectors are colored by phylum. Outer tracks show, for each taxonomic family, stacked counts of database-homologous and database-novel candidates and the number of biomes in which that family was detected. **c**, Species-based rarefaction of candidate sequence families among 18,419 named GTDB species. The curve shows the accumulation of unique families as species are added. Observed richness is 106,764 families, of which 90.5% occur in a single named species. **d**, Family-level biome-private proportions for eight focal phyla, with remaining bacterial and archaeal families grouped as “Other bacteria” and “Other archaea”. Each observation is a GTDB family with at least one scored candidate. Boxes show the median and interquartile range; whiskers extend to 1.5× the interquartile range; outliers are omitted. n values indicate the number of families. Dashed lines mark peptide-weighted domain totals (Bacteria 0.595; Archaea 0.730).

Candidate peptide repertoires exhibited pronounced species-level partitioning. The 18,419 species represented in the analysis collectively encoded 106,764 cAMP families, 90.5% of which were detected in only one species. Among these species-restricted families, 75,145 (77.7%) were biome-private. Species-based rarefaction showed little indication of saturation, with additional species continuing to contribute previously unobserved families (Fig. 3c). These results indicate extensive species-level partitioning of candidate peptide diversity within GIHMsC. Specifically, candidates encoded by CPR bacteria and DPANN archaea showed pronounced biome specificity (Fig. 3d). Across taxonomic families, the median proportion of candidates belonging to biome-private sequence families reached 79.9% in Patescibacteriota and 79.8–83.3% across Nanobdellota, Iainarchaeota, Micrarchaeota and Aenigmatarchaeota, with similarly high proportions in Asgardarchaeota (87.0%), Cyanobacteriota (76.5%) and Bacteroidota (70.9%). For episymbiotic CPR and DPANN lineages, this strong habitat partitioning aligns with their reported site-specific community turnover across isolated ecosystems [50].

### MicroGrad automates dilution for rapid susceptibility tests

To accelerate experimental validation of computationally prioritized AMPs, we developed MicroGrad, a centrifugal microfluidic chip integrating peptide dilution, independent bacterial suspension metering and WST-8 colorimetry to provide an antimicrobial susceptibility readout after 4 h of incubation (within ~4 h from sample loading to visual readout; Fig. 4a, d). Under centrifugal actuation, complementary chamber volumes pair decreasing volumes of peptide solution with increasing volumes of LB medium to generate five twofold concentrations spanning 64 - 4 μmol l□^1^. This volume-compensation design, combined with independent bacterial metering, maintains a consistent nominal inoculum and final assay volume across detection units.

**Figure 4.**
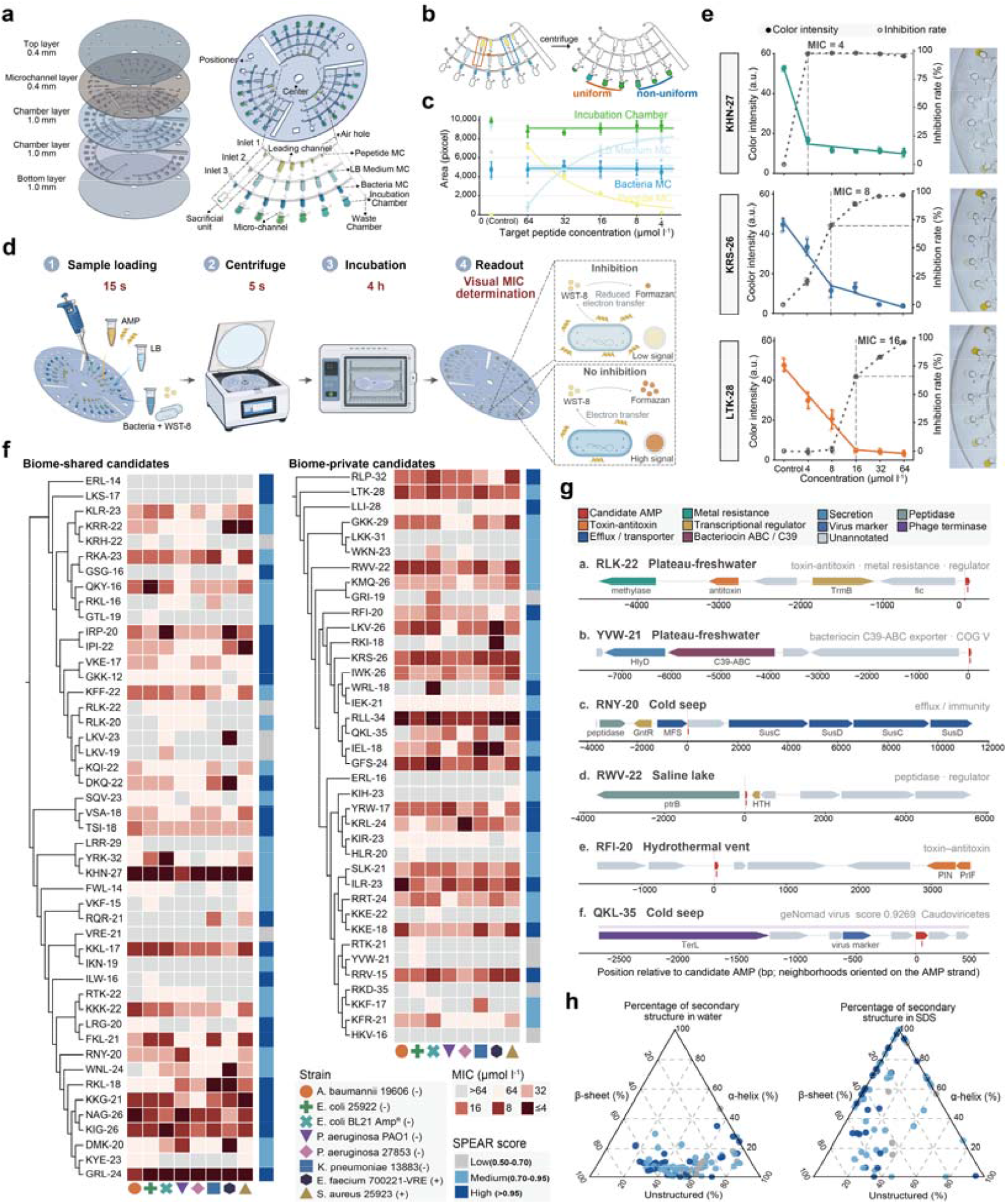
Rapid microfluidic susceptibility testing and characterization of antimicrobial peptide candidates. **a**, Exploded view of the five-layer poly (methyl methacrylate) centrifugal microfluidic chip and schematic of its functional units, comprising three independent inlets, a dam-based flow-splitting and metering structure (Dam-D), and chambers for peptide metering, LB medium compensation, bacterial metering and incubation. Complementary peptide and medium volumes generate the concentration gradient, while independent bacterial metering maintains a consistent nominal inoculum and final assay volume. **b**, Fluidic resistance optimization, showing representative channel configurations and the resulting uniform or non-uniform liquid distribution after centrifugation. Connecting microchannels and local serpentine sections were adjusted to limit overfilling and unintended overflow during liquid transfer. **c**, Liquid-filled areas in the peptide, LB medium and bacterial metering chambers and the incubation chambers across target peptide concentrations. Solid lines represent theoretical values for volume-compensated generation of a twofold concentration gradient. Linear fits for peptide and LB medium allocation yielded R^2^ values of 0.988 and 0.978, respectively. Mean CVs were 3.15% for bacterial metering chambers and 4.89% for incubation chambers. **d**, Chip operation workflow comprising sample loading (15 s), centrifugation (5 s), incubation (4 h) and visual MIC determination. The inset illustrates the WST-8 colorimetric detection principle. Metabolically active bacteria reduce WST-8 to colored formazan, producing a strong colorimetric signal; antimicrobial inhibition reduces formazan formation and signal intensity. **e**, Comparison of on-chip colorimetric readouts and off-chip WST-8 inhibition measurements for KHN-27, KRS-26 and LTK-28. Filled colored symbols and solid lines indicate relative grayscale values extracted from chip images (left axes); open gray symbols and dashed lines indicate inhibition rates calculated in independent off-chip assays (right axes). Corresponding chip images are shown on the right. **f**, MIC heatmaps for 85 candidates sampled across low (0.50-0.70; n = 10), medium (0.70-0.95; n = 43) and high (>0.95; n = 32) SPEAR-score strata. The bacterial panel comprised *Acinetobacter baumannii* ATCC 19606, *Escherichia coli* ATCC 25922, an ampicillin-resistant *E. coli* BL21 strain, *Klebsiella pneumoniae* ATCC 13883, *Pseudomonas aeruginosa* ATCC 27853, *P. aeruginosa* PAO1, *Staphylococcus aureus* ATCC 25923 and vancomycin-resistant *Enterococcus faecium* ATCC 700221. Symbols identify bacterial strains, with − and + denoting Gram-negative and Gram-positive strains, respectively. Candidates are grouped by membership in sequence families detected in multiple biomes (biome-shared) or a single biome (biome-private). Dendrograms were derived from FoldMason multiple-structure alignments. Heatmap colors indicate MICs, and adjacent annotation bars indicate SPEAR-score strata. **g**, Genomic neighborhoods of six representative candidates, with candidate identifiers and source habitats indicated. Arrows represent genes and their orientations; colors indicate predicted functional categories, with candidate AMP genes highlighted in red. Coordinates are relative to the candidate AMP gene, and neighborhoods are oriented consistently with its encoding strand. Selected gene annotations are shown. The predicted viral region containing QKL-35 is annotated with its geNomad virus score and taxonomic assignment. **h**, Secondary-structure compositions measured by circular dichroism spectra in water (left) and in the presence of sodium dodecyl sulfate (SDS; right). Ternary coordinates indicate the proportions of α-helical, β-sheet and unstructured conformations. Each point represents one candidate and is colored by SPEAR-score stratum. MC, medium compensation; WST-8, water-soluble tetrazolium salt-8; LB, Luria-Bertani medium; a.u., arbitrary units; MIC, minimum inhibitory concentration.

Reliable gradient generation requires accurate liquid metering across chambers of different sizes. During centrifugal transfer, rapid liquid delivery from larger to smaller chambers could cause transient local pressure increases, leading to overfilling and unintended overflow into adjacent chambers. To limit this behavior, we adjusted the hydraulic resistance of the connecting microchannels and local serpentine sections (Fig. 4b). In dye experiments, this optimization reduced the mean coefficient of variation (CV) of dye-filled areas across three adjacent culture chambers from 18.17% to 4.88% (n = 6) (Supplementary Fig. 7b, c), ensuring uniform chamber filling.

We next evaluated the metering and mixing performance of the optimized chip using dye solutions (Fig. 4c, Supplementary Fig. 7a). The solutions were loaded sequentially through the three inlets and distributed under centrifugal actuation. Dye allocation to the AMP dosing chambers decreased across the prescribed concentration series, whereas allocation to the LB compensation chambers increased correspondingly. Linear fits for the two allocation series yielded R^2^ values of 0.988 and 0.978, respectively (n = 8). The mean CVs of dye-filled areas in the bacterial metering chambers and final culture chambers were 3.15% and 4.89%, respectively, confirming precise volumetric metering and gradient generation across all detection units.

To assess biological assay performance, we tested three newly identified candidates spanning different potency levels, KHN-27, KRS-26 and LTK-28, and compared their on-chip WST-8 colorimetric responses with inhibition measurements from independent off-chip *microplate* assays (Fig. 4e). WST-8 provides a metabolic readout through formation of a colored formazan product; reduced color development therefore indicates lower bacterial metabolic activity. For all three peptides, increasing peptide concentration reduced the on-chip color signal, with a pronounced transition at a peptide-specific concentration. For KHN-27, the signal decreased from 52.71 to 16.53 arbitrary units (a.u.) at 4 μmol l□^1^. Corresponding decreases were observed for KRS-26 at 8 μmol l□^1^, from 33.28 to 11.39 a.u., and for LTK-28 at 16 μmol l□^1^, from 20.57 to 4.85 a.u. The off-chip inhibition rates at these transition concentrations were 97.58%, 69.82% and 65.57%, respectively (n = 3). These results demonstrate that MicroGrad resolves distinct concentration-dependent inhibitory responses while integrating automated dilution and bacterial metering with a rapid colorimetric readout.

### Top candidate AMPs exhibit potent, broad-spectrum activity

We synthesized and validated 85 candidate AMPs encoded by database-novel smORFs. These peptides were sampled across three SPEAR-score intervals: 10 in the Low group (0.50-0.70), 43 in the Medium group (0.70-0.95) and 32 in the High group (>0.95). For systematic cross-strain profiling, we tested these peptides in broth microdilution assays against eight bacterial strains representing six clinically relevant species: *Acinetobacter baumannii, Escherichia coli, Klebsiella pneumoniae, Pseudomonas aeruginosa, Staphylococcus aureus* and *Enterococcus faecium*. The panel included ampicillin-resistant *E. coli* and vancomycin-resistant *E. faecium*. Overall, 79 of the 85 peptides (92.9%) exhibited antimicrobial activity against at least one strain at MIC ≤64 μmol l□^1^ (Fig. 4f).

Of the tested peptides, 30 (35.3%) met our high-potency criterion of MIC ≤16 μmol l□^1^ against at least four strains or MIC ≤8 μmol l□^1^ against at least two strains. The proportion meeting this criterion increased across the SPEAR-score groups, from 0% in the Low group (0/10) to 34.9% in the Medium group (15/43) and 46.9% in the High group (15/32). Several candidates combined high potency with broad-spectrum activity. Notably, KHN-27, GRL-24 and RLL-34 inhibited all eight strains at MICs ≤8 μmol l□^1^, encompassing both Gram-negative and Gram-positive bacteria and the two antibiotic-resistant strains. Furthermore, genomic-context analysis of validated AMPs revealed diverse functional neighbourhoods, including genes associated with transport, transcriptional regulation and toxin-antitoxin systems (Fig. 4g). YWW-21 occurred near genes encoding a C39 peptidase-containing ABC transporter and HlyD, suggesting a potential connection to peptide processing and export. QKL-35 was located within a phage-associated region containing a terminase large-subunit gene and a viral marker, highlighting a phage-associated genomic context for this AMP.

To examine the conformational and supramolecular properties of the validated candidates, we characterized their secondary structures and self-assembly behavior. Circular dichroism spectroscopy showed that the validated AMPs were predominantly disordered or antiparallel β-sheet-like in water, whereas membrane-mimetic SDS generally increased their α-helical content (Fig. 4h and Supplementary Fig. 8a). However, the SDS-induced increase in helicity was not significantly correlated with strain-specific MICs (Supplementary Fig. 8b-i), consistent with our computational finding that antimicrobial potency is not restricted to a single structural archetype (Supplementary Fig. 4). Dynamic light scattering and atomic force microscopy further revealed concentration-dependent self-assembly in PBS, resolving nanofibres, nanospheres and hook-like nanostructures 8-30 nm in height (Supplementary Fig. 9). Thus, the experimentally validated candidates span diverse conformational transitions and supramolecular states rather than relying on uniform helix induction for activity.

### Potent AMPs disrupt membranes through distinct modes

To investigate the membrane-targeting mechanisms of the potent AMPs prioritized by SPEAR, we used fluorescence assays to compare their activities with the lipopeptide antibiotic Polymyxin B. We examined *Escherichia coli* and *Staphylococcus aureus* as representative Gram-negative and Gram-positive pathogens, respectively. Outer-membrane permeabilization was measured in E. coli using N-phenyl-1-naphthylamine (NPN), which is excluded by an intact outer membrane but becomes fluorescent when membrane disruption exposes it to a hydrophobic lipid environment. Cytoplasmic-membrane depolarization was monitored in both organisms using the membrane-potential-sensitive dye DiSC3(5). The dye accumulates and self-quenches in polarized bacterial membranes; dissipation of the membrane potential releases the dye and increases fluorescence (Fig. 5a-c, Supplementary Fig. 10-12).

**Figure 5.**
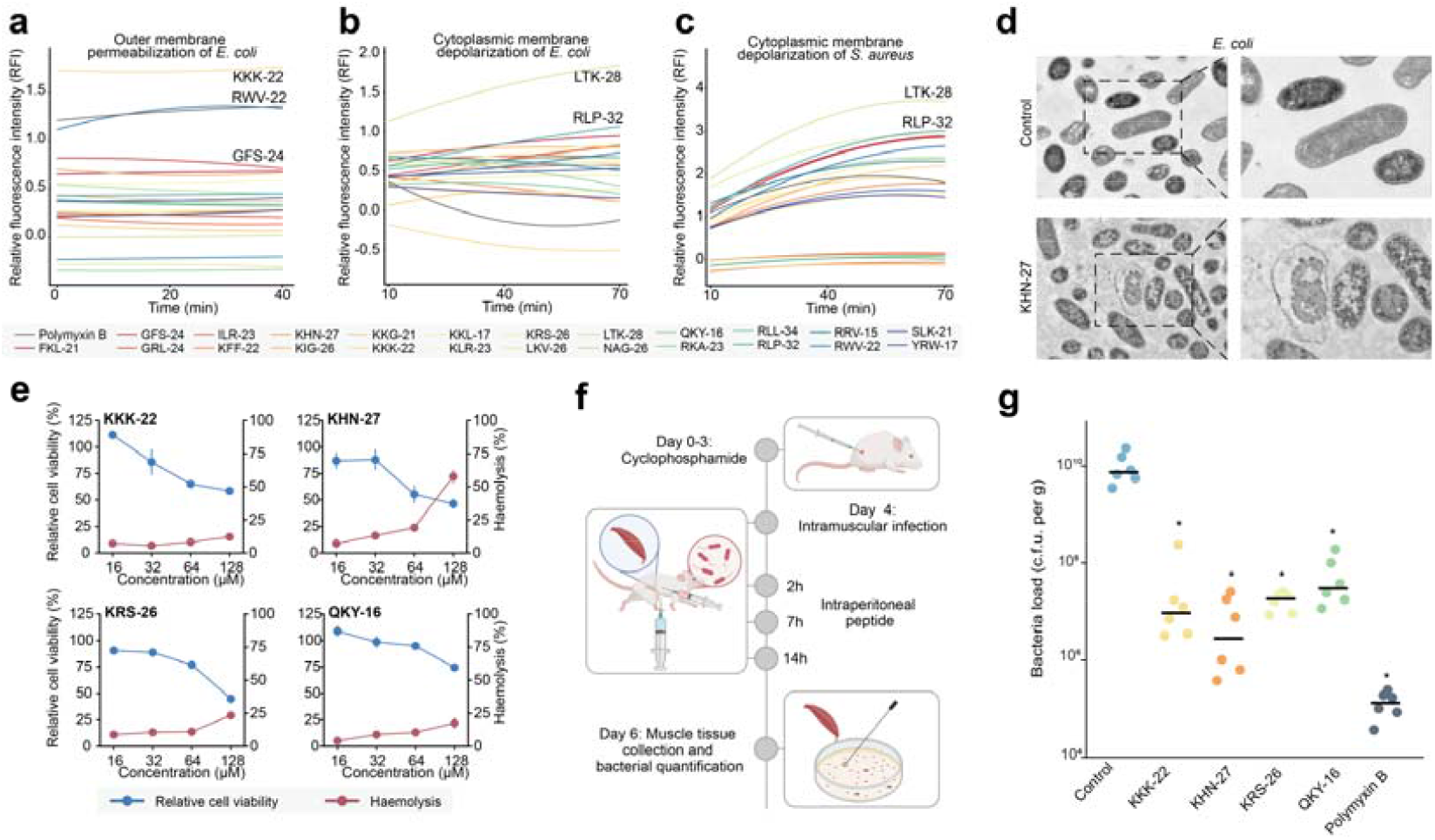
Membrane-disruptive activity, toxicity profiles and therapeutic efficacy of selected AMPs. **a**, Outer-membrane permeabilization of *Escherichia coli* measured using the N-phenyl-1-naphthylamine (NPN) uptake assay. Relative fluorescence was monitored for 40 min after treatment with the indicated peptides at 1× MIC. **b-c**, Cytoplasmic-membrane depolarization of *Escherichia coli* (**b**) and *Staphylococcus aureus* (**c**) measured using the membrane-potential-sensitive dye DiSC3(5). Relative fluorescence was monitored for 60 min after peptide addition at 1× MIC. Increased DiSC3(5) fluorescence indicates dissipation of the transmembrane potential. **d**, Representative transmission electron microscopy images of untreated *Escherichia coli* and cells treated with KHN-27 at 1× MIC. Dashed boxes indicate regions shown at higher magnification on the right. **e**, Cytotoxicity and haemolytic activity of four antimicrobial peptides selected for in vivo validation. Relative cell viability (blue, left y-axis) and hemolysis (red, right y-axis) following exposure to KKK-22, KHN-27, KRS-26 and QKY-16 at concentrations of 16, 32, 64 and 128 μmol l□^1^. **f**, Experimental design of the neutropenic murine deep-thigh infection model. Neutropenia was induced with cyclophosphamide before infection (days 0–3). Mice were infected intramuscularly on day 4. Peptides were administered intraperitoneally at 2, 7 and 14 h after infection. Thigh muscle was collected for bacterial enumeration on day 6 (2 days after infection). Infected controls (PBS) and polymyxin B-treated mice received a single intraperitoneal dose at 12 h after infection (not shown). **g**, Bacterial burdens in infected thigh muscle following treatment with KKK-22, KHN-27, KRS-26, QKY-16 or Polymyxin B. Data are shown as scatter plots with individual mice indicated by dots (n = 6 mice per group). Statistical significance was assessed using one-sided two-sample permutation tests relative to the PBS-treated infected control (*P < 0.01).

Joint analysis of the two fluorescence readouts resolved three membrane-targeting profiles among the potent AMPs: depolarization-dominant, permeabilization-dominant and balanced (Fig. 5a–c). LTK-28 and RLP-32 exemplified the depolarization-dominant profile, inducing strong DiSC□(5) fluorescence with minimal NPN uptake, which indicates preferential disruption of the cytoplasmic membrane over the Gram-negative outer membrane. Conversely, KKK-22 and polymyxin B defined the permeabilization-dominant profile, generating strong NPN uptake with weak endpoint DiSC□(5) signals. Most potent peptides, including FKL-21, YRW-17 and KIG-26, displayed a balanced profile that perturbed both membrane barriers, consistent with their broad-spectrum activity against Gram-negative and Gram-positive pathogens (Fig. 4f). Together, these results show that SPEAR prioritizes both dual-membrane disruptors and peptides with specialized membrane-targeting modes.

PI uptake further distinguished membrane permeabilization from physical loss of cytoplasmic-membrane integrity. At 1× MIC, LTK-28 and ILR-23 produced the strongest PI signals, followed by KHN-27 and RKA-23; their mean fluorescence values exceeded that of Polymyxin B, a potent outer-membrane-permeabilizing antibiotic (Supplementary Fig. 13). In contrast, QKY-16 and RRV-15 elicited weak PI uptake despite strong DiSC□(5) responses, indicating that transmembrane potential collapse can proceed without large pore formation. Furthermore, TEM provided ultrastructural evidence of membrane damage by KHN-27, a representative AMP with a balanced membrane-response profile. Untreated *E. coli* cells retained smooth, continuous envelopes and uniformly electron-dense cytoplasm, whereas KHN-27-treated cells exhibited irregular and disrupted envelopes, membrane detachment, electron-lucent regions and heterogeneous cytoplasmic density (Fig. 5d). Extracellular membranous and debris-like materials further indicated membrane fragmentation and leakage of cellular contents. These ultrastructural changes confirm that potent candidates kill bacteria through physical envelope rupture and cytoplasmic disorganization.

### Low-toxicity AMPs reduce bacterial burdens in mice

We assessed hemolysis in rat red blood cells and cytotoxicity in HEK293T cells for 20 peptides with median MICs ≤32 μmol l□^1^ (Supplementary Fig. 14). Seventeen peptides had HC□□ values >128 μmol l□^1^, and 14 had CC□□ values >32 μmol l□^1^, indicating that most peptides exhibited limited hemolysis at the highest concentration tested and retained >50% cell viability at 32 μmol l□^1^. Applying thresholds of an *Escherichia coli* MIC ≤16 μmol l□^1^ and HC□□ and CC□□ values >64 μmol l□^1^ prioritized KKK-22, KHN-27, KRS-26 and QKY-16 for in vivo evaluation, corresponding to minimum in vitro safety margins of >4 relative to their MICs (Fig. 5e).

We next evaluated the therapeutic efficacy of the four selected peptides in a cyclophosphamide-treated murine thigh infection model (Fig. 5f). Mice were infected intramuscularly with 10□ c.f.u. of *Escherichia coli* and received three intraperitoneal doses of peptides at 10 mg kg□^1^, administered at 2, 7 and 14 h after infection. Control groups received PBS or a single intraperitoneal dose of polymyxin B at 5 mg kg□^1^ at 12 h. Thigh tissues were collected on day 6 for bacterial enumeration. All four peptides significantly reduced thigh bacterial burdens by approximately two to four orders of magnitude compared with PBS-treated controls (Fig. 5g; all P < 0.01, one-sided two-sample permutation tests). Peptide treatment reduced thigh bacterial burdens by approximately 2-4 orders of magnitude relative to PBS-treated infected controls. KHN-27 showed the greatest numerical reduction among the peptide groups, whereas polymyxin B yielded the lowest bacterial burden overall. Serum ELISA measurements further showed lower TNF-α, IL-6 and IL-10 concentrations in all four peptide-treated groups than in PBS-treated infected controls, with the lowest levels observed following polymyxin B treatment (Supplementary Fig. 15). Together with the reduced bacterial burdens, these findings were consistent with attenuation of infection-associated systemic immune activation.

## Discussion

Here, we present MicroSPEAR, an end-to-end framework that combines reliable computational prioritization with rapid, simplified experimental validation for AMP discovery from isolated-habitat metagenomes. Applied to GIHMsC, it yielded experimentally validated antibacterial candidates from sequences underrepresented in existing databases.

GIHMsC highlights the discovery potential of isolated habitat microbiomes. The abundance of candidates without detectable homologues in reference databases and the marked partitioning of candidate families among habitats indicate that these environments contain substantial peptide diversity underrepresented in existing resources. Candidate repertoires in lineages such as CPR bacteria, DPANN archaea and Asgard archaea further extend the taxonomic scope of AMP exploration. They also motivate investigation of whether these peptides contribute to microbial interactions and persistence in their native environments.

SPEAR retained specificity at high recall for low-homology peptides and recovered errors shared by established predictors. External evaluation showed that its scores also ranked peptides by potency despite training on binary labels. However, the contribution of structure-aware representations requires further interpretation. Although ablation analyses support their predictive benefit, the ESMFold-derived structures are not conditioned on the solution or membrane environments in which peptide activity is measured. Comparisons with experimental structural measurements under defined conditions could clarify which features contribute to prediction and how structural uncertainty affects prioritization.

MicroGrad complements this computational capability by reducing the time and liquid-handling effort required for activity testing. On-chip concentration-gradient generation and WST-8 colorimetry enabled MIC determination after 4 h of incubation, with a directly visible readout consistent with off-chip microplate measurements. However, the chip currently requires manual alignment and bonding of its five layers. Standardized manufacturing and preassembled chips could simplify preparation and enable routine multi-strain screening at larger scales.

Together, these computational and experimental capabilities establish a foundation for lab-in-the-loop antibiotic discovery. Although deployed sequentially here, MicroSPEAR is readily adaptable to iterative closed-loop learning, in which model-guided candidate selection informs rapid on-chip susceptibility testing and standardized MIC readouts across active, weakly active and inactive peptides continuously refine potency prediction. More broadly, pairing GIHMsC—which also serves as a genomic resource for mining diverse bioactive peptides and biosynthetic products—with MicroSPEAR provides a generalizable route from Earth’s isolated microbial sequence space to therapeutic leads.

## Methods

### Construction of global isolated-habitat microbiome smORF catalog

#### Large-scale metagenomes

153,243 metagenome-assembled genomes (MAGs) from 4,480 metagenomic samples were collected and downloaded from the NCBI Bioproject Browser (https://www.ncbi.nlm.nih.gov/bioproject), European Bioinformatics Institute BioStudy Browser (https://www.ebi.ac.uk/biostudies/), SPIRE (https://spire.embl.de/) and National Genomics Data Center (https://ngdc.cncb.ac.cn/). The completeness and contamination of MAGs were assessed using CheckM2 (version 1.1.0) [39] with the ‘allmodels’ mode.

#### Taxonomic annotation

The taxonomic annotation of MAGs was conducted using the Genome Taxonomy Database Toolkit (GTDB-Tk v2 [51]) against the GTDB R232 [52] via the ‘classify_wf’ workflow with default parameters. Then, the phylogenetic trees were inferred using pplacer [53], based on maximum-likelihood placement of MAGs onto the GTDB-Tk reference tree.

#### smORF prediction and clustering

Pyrodigal [54] was applied to predict smORFs from the metagenomes, yielding 6,224,518 sequences ranging from 10 to 50 amino acids (AAs) in length. These sequences were further clustered into protein families using CD-HIT [55] with the following parameters: -c 0.5 -n 2 -T 128 -g 1 -s 0.95 –aL 0.95-l 5. This strategy grouped sequences sharing ≥50% sequence identity, with a minimum coverage of 95% for the shorter sequence, resulting in 3,476,289 protein families.

#### Sequence redundancy for search and scoring

The 6,224,518 predicted smORFs constituted the GIHMsC catalog used for habitat and family counts. For homology searches, identical sequences were collapsed to 5,297,597 unique peptides. DIAMOND hits were then mapped back to all 6,224,518 catalog entries, yielding 2,696,886 homologous smORFs (43.3%) and 3,527,632 database-novel smORFs (56.7%). For SPEAR scoring, the search set was further reduced to 4,738,802 sequences by excluding sequences that failed ESMFold. Family counts (3,476,289 families at 50% identity) were computed on the full catalog and were not used as the SPEAR input.

#### Large-scale homolog search against databases

To assess novelty, we compiled all proteins from SmProt2 [56] (*n* = 449,335), UniRef90 [57] (*n* = 153,742,194), Refseq-bacteria (*n* = 384,470,870), Refseq-archaea (*n* = 4,656,312) [58], sORFdb (*n* = 34,007,166) [59] and GMSC [60] (*n* = 964,970,496). All nonredundant sequences from the catalog (*n* = 5,297,597) were searched against the combined datasets (*n* = 1,542,296,373) using DIAMOND in ‘-more-sensitive’ mode, retaining hits with *E value* < 10^−5^. In total, 2,696,886 (43.3%) smORFs showed homology to known proteins.

### Dataset construction

We assembled three datasets for model training and evaluation.

#### AMP prediction module training dataset

To obtain a diverse AMP classification dataset with broad coverage of different data sources, we integrated sequences from curated AMP databases, a general-purpose protein database, a previously published benchmark dataset, and experimentally validated literature datasets. Specifically, the dataset was assembled from four sources: (i) AMPs from DBAASP [40] and DRAMP [41] (primary positive dataset); (ii) small proteins without annotated antimicrobial activity obtained from UniProt (primary negative dataset) [42]; (iii) the HMD-AMP training dataset [43], and (iv) 303 experimentally validated peptides collected from six published studies [8, 61–65]. For the UniProt-derived negative dataset, proteins annotated with a cytoplasmic subcellular location were retained, whereas entries associated with “antimicrobial”, “antibiotic”, “antiviral”, “antifungal”, “antibacterial”, “bacteriocin”, “effector”, or “excreted” were excluded. Across all sources, sequences containing non-canonical amino acids, D-amino acids, or chemical modifications were removed, and only sequences of 10-50 amino acids were retained. To further reduce redundancy and increase sequence diversity, the combined dataset was clustered using CD-HIT at a 75% sequence-identity threshold, with one representative sequence retained from each cluster. The final dataset contained 13,652 sequences, comprising 6,169 AMPs and 7,483 non-AMPs.

#### Baseline consensus-failure test set

To determine whether the evaluated models could correct errors shared by established AMP predictors, we constructed a baseline consensus-failure (BCF) test set from the training data. Macrel [12], AMPlify [15], and APEX 1.1 [46] were selected as fixed baselines because they are widely used AMP predictors. Their released pretrained models and fixed inference settings were used without retraining or optimization on the classification dataset. Because APEX 1.1 predicts strain-specific minimum inhibitory concentration (MIC) values rather than binary labels, we converted its outputs using the following rule. A sequence was classified as an AMP if APEX 1.1 predicted an MIC below 128 µM for at least one strain; otherwise, it was classified as a non-AMP. Each sequence was evaluated independently by the three baseline models, and the resulting binary predictions were compared with the ground-truth label. Sequences misclassified by at least two of the three baseline models were assigned to the BCF test set. The remaining sequences constituted the training set for the models evaluated in this study. No sequence in the test set was used for model training. This failure-conditioned partition enabled direct assessment of whether an evaluated model could correctly classify sequences for which most fixed baseline predictors failed.

#### External test dataset

Model performance was evaluated on five non-overlapping peptide collections: 79 experimentally validated peptides from Santos-Júnior et al. [9], 39 peptides from Torres et al. [49], 54 peptides from Guan et al. [48], 48 food-derived peptides from our in-house dataset, and 113 peptides from Jiang et al. [47]. We selected experimentally characterized peptides from individual studies rather than public AMP databases for three reasons. First, to the best of our knowledge, these peptides had not been used to train the baseline models, reducing the risk of test-set leakage. Second, MIC measurements for 51.7 % of the peptides were generated by a single laboratory (the de la Fuente-Nunez laboratory), helping to limit interlaboratory variability in the activity data. Third, these peptides had already been prioritized as AMP-like candidates by computational methods, providing a challenging setting for SPEAR and the baseline models. The combined test set comprised 333 unique sequences. We then classified the peptides as non-AMPs (n = 144; MIC >64 μmol l□^1^ against all strains with reported measurements, or no exact MIC recorded), weak-AMPs (n = 47; MIC ≤64 μmol l□^1^ against only one strain), or AMPs (n = 142; MIC ≤64 μmol l□^1^ against at least two strains). For the 220 peptides with experimental MICs, measurements were available against 6 to 11 strains drawn from six clinically relevant bacterial pathogens: Acinetobacter baumannii, Escherichia coli, Klebsiella pneumoniae, Pseudomonas aeruginosa, Staphylococcus aureus and Enterococcus faecium.

### Physicochemical property calculation

The physicochemical properties of AMPs were calculated using the *ProteinAnalysis* module in *Biopython* and the Python package *Peptides* [66]. *Biopython* was used to calculate molecular weight, peptide length, aromaticity and the grand average of hydropathicity (GRAVY). *Peptides* was used to calculate the instability index, isoelectric point, net charge, Boman index, hydrophobicity, hydrophobic moment and aliphatic index. The Boman index estimates the protein-binding potential of a peptide based on the average solubility free energy of its constituent amino acids [67]. These descriptors collectively characterize peptide size, charge, stability, hydrophobicity, amphipathicity, and potential protein-binding capacity.

### Model architecture

SPEAR comprised a dual-view representation module and an ensemble classifier. The representation module encoded each peptide from both its primary amino acid sequence and its predicted three-dimensional structure. The resulting sequence and structure-aware representations were concatenated and supplied to two complementary tree-based classifiers, whose output probabilities were averaged to obtain the final antimicrobial peptide score.

#### Representation extraction

For a peptide *S* = (*a*_1’_,’*a*_2’_,’…’,’*a*_*L*_) of length *L*, where *a*_*i*_ is the amino acid at position *i*. Two complementary representations were extracted for each peptide. For the sequence view, the amino acid sequence was provided to the pretrained 650-million-parameter ESM-2 protein language model. The final hidden layer produced residue-level contextual embeddings. Given the scale of the peptide dataset and the computational cost of structure prediction, the three-dimensional structure of each peptide was inferred using ESMFold [44]. The predicted structure 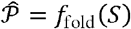 was restricted to chain A and converted into the Foldseek [68] 3Di sequence *Q* = (*q*_1’_,’q_2’_,’…’,’*q*_*L*_), where *q*_*i*_ describes the local tertiary interaction environment at residue *i*. The amino acid and 3Di symbols at corresponding positions were then combined into structure-aware sequence *U* = ((*a*_1’_,’*q*_1_)’,’…’,’(*a*_*L’*_,’*q*_*L*))_, which was subsequently processed by a pretrained SaProt model, producing 695 residue-level structure-aware embeddings. A fixed-dimensional peptide representation 696 was obtained separately from each model by mean-pooling the final-layer residue 697 embeddings. Padding and model-specific beginning- and end-of-sequence tokens 698 were excluded from the pooling operation. The sequence and structure-aware 699 embeddings were further concatenated to form the final feature vector:

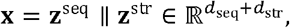

where ║ denotes vector concatenation.

#### Ensemble classification

The concatenated representations were classified using an ensemble comprising a random forest (RF) and a histogram-based gradient-boosting machine (HistGBM) classifier. The random forest contained 1,000 decision trees, whereas the gradient-boosting classifier was trained for a maximum of 100 boosting iterations. The two classifiers were trained independently on the same feature vectors. Peptides annotated as antimicrobial were assigned *y* =1, and all remaining peptides were assigned *y*=0. Each decision tree was fitted to a bootstrap sample of the training data, and a random subset of features was considered at each node. For HistGBM classifier, continuous features were discretized into integer-valued bins before training. At each iteration, histogram-aggregated gradients and Hessians were used to fit a regression tree to the current errors, and its leaf values were added to the accumulated logit. The final SPEAR score was defined as the unweighted arithmetic mean of the two positive-class probabilities. Random seeds were fixed at 42 for both classifiers to ensure reproducible model fitting.

### Evaluation metrics

#### Classification performance and error correction

To evaluate the classification performance of SPEAR and competing models, we used specificity, recall, F1-score and Matthews correlation coefficient (MCC), calculated as follows:

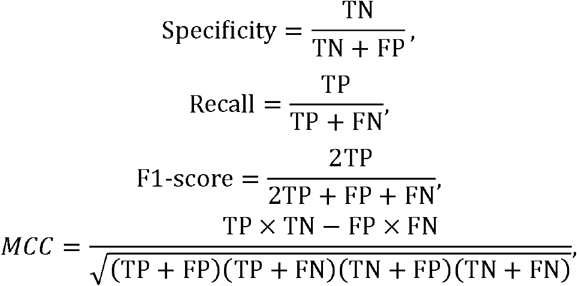

where TP, TN, FP and FN denote the numbers of true positives, true negatives, false positives and false negatives, respectively. To assess the ability to reject negative samples while retaining high recall, we used Specificity@Recall0.90:

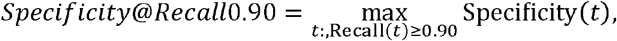

where *t* denotes the model decision threshold. Error correction rates (ECRs) were evaluated on the consensus-failure test set:

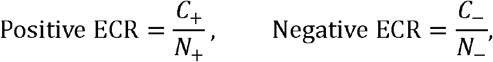

Where *N*_+_ and *N*_-_ denote the numbers of positive and negative samples in this test set, and *C*_+_ and *C*_-_ denote the corresponding numbers correctly classified by the evaluated model.

#### Probability calibration and prediction error

To assess the agreement between predicted AMP probabilities and experimental labels, we used expected calibration error (ECE) and the Brier score. ECE measures the discrepancy between mean predicted probabilities and observed positive-class frequencies across ten equal-width probability bins, whereas the Brier-score measures the mean squared error of probabilistic predictions:

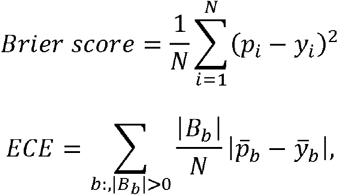

where *N* denotes the number of evaluated peptides, *p*_*i*_734 is the predicted AMP probability and *y*_*i*_ is the reference label. *B*_*b*_ denotes the set of peptides in probability bin *b*, and 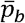 and 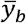 denote its mean predicted probability and observed positive-class fraction, respectively. Lower values indicate better calibration for ECE and lower probabilistic prediction error for the Brier score.

#### Spectrum-adjusted antimicrobial potency

To integrate antimicrobial potency and spectrum across tested bacterial species, we calculated a spectrum-adjusted potency index (SAPI). MIC values were converted into normalized activity scores and averaged first across strains within each species and then equally across tested species:

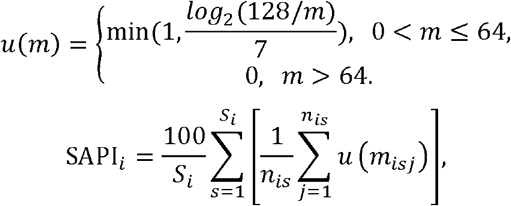

where *m* denotes the numerical MIC value expressed in μmol l □^1^,*S*_*i*_ is the number of species tested for peptide, *i, n*_*is*_ is the number of strains tested within species *S*, and *m*_*isj*_ is the MIC against strain *j* of species *S*. MIC values ≤1 μmol 1□^1^ were assigned an activity score of 1, an MIC of 64 μmol l□^1^ was assigned a score of 1/7, and MIC values >64 μmol l□^1^ contributed zero activity. Untested species and strains were excluded from the corresponding averages. SAPI ranges from 0 to 100, with higher values indicating stronger aggregate antimicrobial activity across the tested species.

### Functional annotation of genomic contexts

Genes surrounding candidate AMP-encoding smORFs were functionally annotated using eggNOG-mapper (v2.1.13), with DIAMOND searches against the database in ultra-sensitive mode (-m diamond --sensmode ultra-sensitive). DefenseFinder (v3.1.0) was used to identify antiphage defense systems and their constituent genes. The corresponding MAGs were analyzed using the geNomad end-to-end workflow (v1.11.2) to identify proviral regions and obtain their virus scores. These annotations were mapped to genomic coordinates to characterize neighboring gene functions and the positions of candidate AMP loci relative to defense systems and proviral regions. Default parameters were used unless otherwise specified.

### Peptide synthesis

Peptides were commercially synthesized by GenScript Biotech Co., Ltd. using solid-phase peptide synthesis (SPPS). The identity of each synthesized peptide was verified by mass spectrometry by comparing the experimentally measured molecular mass with its corresponding theoretical value.

### Chip design and fabrication

The chip was designed in AutoCAD (Autodesk) and fabricated by CO□ laser cutting of poly (methyl methacrylate) (PMMA) sheets. The five layers were cleaned with nitrogen gas and assembled using double-sided adhesive films, with absorbent paper pads inserted into the designated chambers before final assembly. The assembled chip was hot-pressed at 50 °C for 10 min to strengthen interlayer bonding.

### Bacterial strains and growth conditions

The Gram-negative panel comprised Acinetobacter baumannii ATCC 19606, Escherichia coli ATCC 25922, an ampicillin-resistant laboratory derivative of E. coli BL21, Klebsiella pneumoniae ATCC 13883, Pseudomonas aeruginosa ATCC 27853, and P. aeruginosa PAO1. The Gram-positive panel comprised Staphylococcus aureus ATCC 25923 and vancomycin-resistant Enterococcus faecium ATCC 700221. E. coli BL21 and E. faecium ATCC 700221 were included as antibiotic-resistant strains, exhibiting resistance to ampicillin and vancomycin, respectively. In all experiments, all strains except *E. faecium* ATCC 700221 were cultured overnight in LB medium at 37 °C, whereas *E. faecium* ATCC 700221 was cultured overnight in TSB under the same conditions. Following cultivation, each bacterial suspension was adjusted to a 0.5 McFarland standard, corresponding to approximately 1.5 × 10^8^ colony-forming units (c.f.u.) mL^-1^, and subsequently diluted 100-fold in the corresponding growth medium to approximately 1.5 × 10^6^ c.f.u. mL^-1^. They were further incubated at 37 °C to mid-logarithmic phase.

### MIC assay

#### On-chip antimicrobial susceptibility testing

The antimicrobial susceptibility testing capability of the MicroGrad chip was evaluated using three antimicrobial peptides, KHN-27, KRS-26 and LTK-28, against Escherichia coli ATCC 25922. Bacteria were cultured to the exponential growth phase and diluted in LB medium to the required inoculum density. Peptide solution, LB medium and bacterial suspension were introduced through their respective chip inlets and subjected to centrifugal actuation for automated metering, distribution and mixing. Complementary volumes in the peptide metering chambers and LB compensation chambers generated a five-step twofold dilution series with final peptide concentrations of 64, 32, 16, 8 and 4 μmol l□^1^. A peptide-free bacterial growth control was included. WST-8 reagent was added to the reaction mixture before incubation as a colorimetric indicator of bacterial metabolic activity. Following liquid distribution, the chip was sealed with PCR sealing film and incubated at 37 °C for 4 h. On-chip MICs were read directly by visual inspection of color development across the concentration series. The endpoint was identified as the lowest peptide concentration on the inhibited, weakly colored side of the transition from pronounced to suppressed color development.

Images of the culture chambers were also acquired for quantitative analysis in ImageJ. Original images were converted to 8-bit grayscale using weighted RGB conversion. Circular regions of interest (ROIs) of identical size were placed within the reaction liquid in each chamber, excluding chamber walls, white circular openings, bubbles and prominent reflections. Mean gray values were measured for the sample, corresponding image background and bacterial growth control. Background-corrected color intensity was calculated by subtracting the sample mean gray value from the corresponding background mean gray value, such that higher values indicated stronger color development. Intensities were expressed in arbitrary units without normalization to the bacterial growth control. For each independent replicate, the growth-control intensity was calculated by subtracting its mean gray value from the average background gray value across the five peptide concentrations.

Corrected color intensity was plotted against peptide concentration to identify the colorimetric breakpoint, defined here as the transition from high to low color intensity with increasing peptide concentration. The lowest tested concentration on the low-intensity side of this transition was assigned as the image-derived MIC and compared with the visual readout. Each assay was performed in three independent replicates, and color intensities are presented as the mean ± standard deviation across replicates. On-chip MICs were compared with the corresponding off-chip growth-inhibition profiles to assess consistency between the two assays.

#### Off-chip antimicrobial susceptibility testing

Activity profiling of the 85 synthesized candidate peptides across the eight-strain bacterial panel was performed in 384-well microplates. Antimicrobial activity was initially assessed by measuring bacterial growth inhibition at a final peptide concentration of 64 μmol l□^1^. Diluted bacterial suspensions were mixed with peptide working solutions prepared at 128 μmol l□^1^ at a 1:1 (v/v) ratio in 384-well microplates, yielding a final bacterial density of approximately 7.5 × 10□ c.f.u. mL□^1^. Bacterial growth was quantified by measuring absorbance using a microplate reader. The inhibition rate was calculated as:

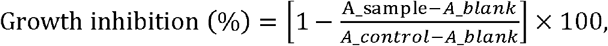

where *A_sample*, *A_control* and *A_blank* denote the absorbance of the peptide-treated bacterial sample, untreated bacterial growth control and blank control, respectively. Peptide solutions were mixed with bacterial suspensions to obtain final peptide concentrations of 64, 32, 16, 8 and 4 μmol l□^1^ under the same assay conditions. Each concentration was tested in three technical replicate wells, with the corresponding growth and blank controls.

### Cytotoxicity assay

The cytotoxicity of the peptides toward human embryonic kidney 293T (HEK293T) cells was assessed using the MTT assay. HEK293T cells were maintained in high-glucose Dulbecco’s modified Eagle’s medium (DMEM) supplemented with 10% fetal bovine serum and 1% penicillin-streptomycin at 37 °C in a humidified atmosphere containing 5% CO□. Cells were seeded into 96-well plates at a density of 5,000 cells per well by dispensing 100 μL of a 50,000 cells/mL suspension. After cell attachment, the culture medium was removed and replaced with fresh complete medium containing the peptides at final concentrations of 16, 32, 64, or 128 μmol l□^1^. Untreated cells served as the viability control, whereas wells containing culture medium without cells served as background blanks. Cells were incubated with the peptides for 24 h. Following peptide treatment, the culture medium was removed, and 100 μL of 0.5 mg mL^-1^ MTT prepared in phenol red-free medium was added to each well. The plates were incubated for an additional 4 h at 37 °C to allow the formation of insoluble purple formazan crystals. The MTT solution was then carefully removed, and the formazan crystals were dissolved in acidified isopropanol containing 0.04 mol L^-1^ HCl. After complete dissolution, the absorbance was measured at 570 nm. Cell viability was calculated as follows:

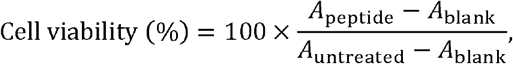

Where *A*_peptide_, *A*_untreated_, and *A*_blank_ represent the absorbance values of peptide-treated cells, untreated cells, and cell-free background wells, respectively.

### Hemolysis assay

The hemolytic activity of the peptides was evaluated using commercially obtained heparinized rat erythrocytes. Erythrocytes were washed four times with phosphate-buffered saline (PBS, pH 7.4) by centrifugation at 800 g for 10 min, with the supernatant carefully discarded after each wash. The washed erythrocytes were subsequently diluted 200-fold in PBS. For the assay, 75 μL of the erythrocyte suspension was gently mixed with 75 μL of peptide solution in a 96-well plate to achieve final peptide concentrations of 16, 32, 64, and 128 μmol l□^1^. Erythrocytes incubated with PBS served as the negative control, representing 0% hemolysis, whereas erythrocytes treated with 1% (v/v) sodium dodecyl sulfate (SDS) in PBS served as the positive control, representing 100% hemolysis. The plates were incubated at room temperature for 4 h. Following incubation, the plates were centrifuged at 1,300 g for 10 min to sediment intact erythrocytes and cellular debris. Subsequently, 100 μL of the supernatant from each well was transferred to a new 96-well plate, and the absorbance was measured at 405 nm. The percentage of hemolysis was calculated as follows:

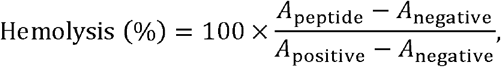

where *A*_peptide_, *A*_negative_, and *A*_positive_ represent the absorbance values of the peptide-treated, PBS-treated, and SDS-treated samples, respectively. Peripheral wells of the plate were not used to minimize edge effects, and all liquid-handling steps were performed gently to avoid mechanically induced erythrocyte lysis.

### Circular dichroism spectroscopy

The secondary structures of the antimicrobial peptides under different environmental conditions were characterized using circular dichroism (CD) spectroscopy. Each peptide was prepared at a final concentration of 50 μmol l□^1^ in either sterile water or an aqueous solution containing 10 mM sodium dodecyl sulfate (SDS). The SDS-containing condition was used to provide a membrane-mimetic hydrophobic environment and assess potential peptide conformational changes between different microenvironments. CD spectra were recorded at 25 °C using a quartz cuvette with a path length of 1 mm. Spectra were collected over a wavelength range of 190~260 nm at a scanning rate of 50 nm min^-1^ and a bandwidth of 0.5 nm. Three consecutive scans were acquired for each sample and averaged. Before sample measurements, baseline spectra were recorded using sterile water or 10 mM SDS, as appropriate, and the corresponding solvent background was subtracted from each peptide spectrum.

### Outer-membrane permeability assay

The effect of antimicrobial peptides on bacterial outer-membrane permeability was evaluated using an N-phenyl-1-naphthylamine (NPN) uptake assay. Following cultivation at 37 °C, bacterial cells were harvested, washed with a buffer containing 5 mM HEPES and 5 mM glucose, and resuspended in the same buffer. The bacterial suspension was adjusted to an OD600 of 0.13. A 100 μL aliquot of the bacterial suspension was mixed with NPN to obtain a final assay concentration of approximately 10 μmol l□^1^, followed by the addition of 100 μL of antimicrobial peptide solution. Changes in fluorescence intensity were continuously monitored for 40 min using a microplate reader with excitation and emission wavelengths of 350 and 420 nm, respectively. Polymyxin B and the corresponding vehicle were used as the positive and negative controls, respectively.

### Cytoplasmic membrane depolarization assay

Antimicrobial peptide-induced cytoplasmic membrane depolarization was evaluated using 3,3′-dipropylthiadicarbocyanine iodides □DiSC3(5)]. Following cultivation, bacterial cells were harvested by centrifugation, washed with a buffer containing 5 mM HEPES and 20 mM glucose (pH 7.4), and resuspended in the same buffer. The bacterial suspension was adjusted to an OD600 of 0.05. DiSC3(5) was added to a 100 μL aliquot of the bacterial suspension to obtain a final assay concentration of 0.4 μmol l□^1^, followed by incubation for 2 h. Subsequently, 100 μL of antimicrobial peptide solution was added, and changes in fluorescence intensity were continuously monitored for 1 h using a microplate reader with excitation and emission wavelengths of 650 and 675 nm, respectively. Polymyxin B and the corresponding vehicle were used as the positive and negative controls, respectively.

### Cytoplasmic membrane integrity assay

The effect of antimicrobial peptides on bacterial cytoplasmic membrane integrity was evaluated using a propidium iodide (PI) uptake assay. A 2 mL aliquot of bacterial culture was centrifuged at 8,000 rpm for 10 min at 4 °C to collect the cells, which were subsequently washed with 0.9% NaCl. The cells were resuspended in 0.9% NaCl, and the bacterial suspension was adjusted to an OD600 of 0.1. A 100 μL aliquot of the bacterial suspension was mixed with 100 μL of antimicrobial peptide solution and incubated for 30 min. PI was then added to a final concentration of 2.5 μg mL^-1^, followed by incubation at 4 °C for 15 min in the dark. After incubation, the samples were centrifuged at 10,000 g for 10 min to remove unbound PI and residual peptide. The supernatant was discarded, and the cell pellet was resuspended in 350 μL of 0.9% NaCl. Changes in fluorescence intensity were continuously monitored for 20 min using a microplate reader with excitation and emission wavelengths of 535 and 617 nm, respectively.

### TEM measurement

Bacterial cells were harvested by centrifugation at 8,000 × g for 10 min. After removal of the supernatant, the bacterial pellets were collected and resuspended in electron microscopy fixative. The samples were fixed overnight and subsequently subjected to conventional TEM sample preparation. Briefly, the fixed samples were washed with phosphate buffer and post-fixed with osmium tetroxide. The samples were then dehydrated through a graded ethanol series, followed by resin infiltration and embedding. After resin polymerization, ultrathin sections were prepared using an ultramicrotome and stained with uranyl acetate and lead citrate. The ultrastructural morphology of the bacterial cells was finally examined using transmission electron microscopy (TEM), and representative images were acquired.

### Thigh infection mouse model

#### Establishment and treatment of mouse model

Six-to eight-week-old female ICR mice were purchased from SPF (Beijing) Biotechnology Co., Ltd. (Beijing, China) and acclimatized for 1 week before experimentation. The mice were divided into six groups, with six animals per group, including an infection control group, four antimicrobial peptide treatment groups, and an antibiotic treatment group. Neutropenia was induced by intraperitoneal administration of cyclophosphamide at 150 mg kg^-1^ on day 1 (3 days before infection) and 100 mg kg^-1^ on day 3 (1 day before infection). Day 0 was defined as the first day of the experiment, and mice were infected on day 4. *Escherichia coli* ATCC 25922 was cultured in Luria–Bertani broth at 37°C to the exponential growth phase. Bacterial cells were collected, washed twice with sterile phosphate-buffered saline (PBS), and resuspended in PBS at 1 × 10^6^ c.f.u. mL^-1^. Each mouse was inoculated intramuscularly in the right thigh with 100 μL of the bacterial suspension, corresponding to an infection dose of 1 × 10^5^ c.f.u. per mouse. Mice in the peptide treatment groups received intraperitoneal injections of the indicated antimicrobial peptide at 2, 7, and 14 h after infection, with each dose administered at 10 mg kg^-1^. Peptide powders were freshly dissolved in sterile PBS before use whenever possible. Mice in the antibiotic treatment group received a single intraperitoneal injection of Polymyxin B at 5 mg kg^-1^ at 12 h after infection. The infection control group received an equivalent volume of sterile PBS at 12 h after infection. All mice were euthanized on day 6 (2 days after infection). The right thigh muscle was aseptically collected for bacterial burden determination, and blood was collected for serum analysis. All animal experimental protocols were reviewed and approved by the Laboratory Animal Welfare & Ethics Committee of the Bestcell Model Biological Center (approval no. BSMS-IACUC-2026-09-23B).

#### Determination of bacterial burden of mouse model

Following euthanasia, the external surface of each mouse was disinfected with 75% ethanol for 5 min. The entire infected right thigh muscle was aseptically excised and weighed. Sterile glycerol was then added at a tissue mass-to-liquid volume ratio of 1 g:1 mL, and the tissue was thoroughly homogenized. The homogenates were serially diluted in sterile PBS, and appropriate dilutions were plated onto eosin methylene blue agar for colony enumeration. Bacterial burden was calculated by correcting for the dilution factor and plated volume and was normalized to the initial tissue mass, with results expressed as CFU g□^1^ of thigh muscle.

#### Serum cytokine measurement by ELISA

Blood samples were centrifuged at 3,000 rpm for 10 min, and the serum fraction was collected. Serum concentrations of tumor necrosis factor-α (TNF-α), interleukin-6 (IL-6), and interleukin-10 (IL-10) were measured using commercial mouse ELISA kits (Jiangsu Meimian Industrial Co., Ltd., Jiangsu, China) according to the manufacturer’s instructions.

## Data availability

The metagenomes used in this study are described in the Supplementary Information and are available from the original sources. The SPEAR-scored candidate AMP catalogue from GIHMsC and the independent external AMP test set are available via Zenodo at https://zenodo.org/records/23904244. All other data generated in this study are available from the corresponding author on reasonable request.

## Code availability

The source codes for SPEAR, including training scripts, and inference scripts, are available at https://github.com/HUST-NingKang-Lab/SPEAR. Peptide secondary structure composition was estimated from circular dichroism spectra using the BeStSel server (https://bestsel.elte.hu/). Peptide structures were visualized using Mol* Viewer (https://molstar.org/viewer/), and phylogenetic trees were visualized using Interactive Tree Of Life (iTOL; https://itol.embl.de/).

## Acknowledgements

This work was partially supported by the National Key R&D Program of China (Grant No. 2023YFA1800900, 2021YFA0910500 and 2018YFC0910502) and the National Natural Science Foundation of China (22574058, 32571640). We thank Dr Célio Dias Santos-Júnior for providing the sequences of peptides tested in vitro in the study “Discovery of antimicrobial peptides in the global microbiome with machine learning”.

During the preparation of this manuscript, the authors used ChatGPT (OpenAI) to improve the readability and language of the text. All AI-assisted edits were reviewed and revised by the authors, who take full responsibility for the final content of the manuscript.

## Author contributions

Z.K., H.Z., K.N. and B.L. conceived the research; Z.K. and H.Z. collected the dataset;

Z.K. implemented the main algorithm; Q.Z. and P.C. designed the microfluidic chip;

Z.K. and Q.Z. designed the experiments; Q.Z., J.L., K.L., K.Z., Y.Z., Y.W. and P.C. performed the experiments; Z.K., Q.Z., H.Z., P.C. and K.N. conducted the analysis; Z.K., Q.Z. and H.Z. wrote the paper; and B.L. and K.N. supervised the project. All authors read and approved the paper.

