## supplemental tables and figures for "Accelerating antibiotic discovery from Earth’s isolated habitats through protein language models and microfluidics"

### List of Supplementary Notes

#### 1. Metagenome data collection and processing

#### 2. Reference AMP coverage and cross-model predictions

#### 3. Structural diversity of reference antimicrobial peptides and its relationships with sequence physicochemical properties and antimicrobial potency

#### 4. Peptide representation evaluation

#### 5. Model ablation study

#### 6. Validation and optimization of liquid distribution in the MicroGrad microfluidic chip

#### 7. Properties of experimentally validated peptides

#### 8. Cytotoxicity and haemolysis of the validated AMPs

#### 9. Serum cytokine responses

### List of Supplementary Figures

#### Supplementary Figure 1 | Quality assessment and taxonomic composition of MAGs recovered from globally distributed isolated microbial habitats

#### Supplementary Figure 2 | Sequence identity to reference AMPs, overlap among predictor candidate sets and cross-model prediction disagreement across smORF families

#### Supplementary Figure 3 | Contact-map-based clustering resolves nine AMP structural archetypes

#### Supplementary Figure 4 | Contact-map-defined AMP structural archetypes are associated with physicochemical properties

#### Supplementary Figure 5 | SPEAR retains complementary AMP-label and structural-archetype information in its fused embedding

#### Supplementary Figure 6 | Ablation of protein language-model representations and classifiers under a 30% sequence-identity constraint

#### Supplementary Figure 7 | Dye-based characterization of liquid distribution and hydraulic resistance optimization in the MicroGrad chip

#### Supplementary Figure 8 | Secondary-structure compositions of all synthesized peptides in water and sodium dodecyl sulfate (SDS) and their associations with strain-specific antimicrobial activity

#### Supplementary Figure 9 | Concentration-dependent aggregation and self-assembly of selected AMPs

#### Supplementary Figure 10 | Outer-membrane permeabilization of Escherichia coli measured using the N-phenyl-1-naphthylamine (NPN) uptake assay

#### Supplementary Figure 11 | Cytoplasmic-membrane depolarization of Escherichia coli measured using the membrane-potential-sensitive dye DiSC3(5)

#### Supplementary Figure 12 | Cytoplasmic-membrane depolarization of Staphylococcus aureus measured using the membrane-potential-sensitive dye DiSC3(5)

#### Supplementary Figure 13 | PI uptake by Escherichia coli

#### Supplementary Figure 14 | Dose-dependent cytotoxicity and haemolysis of the tested peptides

#### Supplementary Figure 15 | Serum cytokine levels following peptide or antibiotic treatment in a mouse muscle infection model

### List of Supplementary Tables

#### Supplementary Table 1 | Habitat classifications of MAG source studies

#### Supplementary Table 2 | Environmental-feature metadata of the 15 isolated habitats

#### Supplementary Table 3 | Summary statistics for AMP structural archetypes

#### Supplementary Table 4 | Benchmark of AMP and non-AMP representation methods

#### Supplementary Table 5 | Benchmark of AMP structure representations

### 1. Metagenome data collection and processing

We collected publicly available metagenome-assembled genomes (MAGs) from studies covering 15 isolated habitat types and compiled sample-level metadata from the corresponding publications and associated source data. Environmental isolation was described using a standardized framework (Supplementary Table 1), and each habitat was characterized by temperature, salinity, UV radiation, water availability, pH, metal concentration, nutrient availability, pressure and oxygen availability (Supplementary Table 2).

All downloaded MAGs were assessed using CheckM2, and genomes with estimated completeness ≥50% and contamination ≤5% were retained for downstream analyses. This filtering retained 119,107 of the 153,243 collected MAGs (77.7%). Across the 15 habitats, median estimated contamination remained below approximately 2%, whereas completeness distributions varied, with higher median values in glacier, acid mine drainage and cave MAGs (Supplementary Fig. 1a, b). Assembly contiguity, as measured by contig N50, varied substantially both within and across habitats (Supplementary Fig. 1f). Additional genome characteristics, including assembly size, GC content, coding density, average gene length and the number of coding sequences, are summarized in Supplementary Fig. 1c–h.

The collected MAGs exhibited distinct phylum-level taxonomic profiles across habitats (Supplementary Fig. 1i). Glacier freshwater, plateau freshwater and saline lake MAG collections shared Pseudomonadota, Bacteroidota and Actinomycetota as major constituents. Within this group, Patescibacteriota accounted for a larger proportion of glacier freshwater MAGs, whereas Bacteroidota contributed more prominently to plateau freshwater and saline lake collections. Both hot springs and hydrothermal vents contained appreciable proportions of Thermoproteota, but their bacterial profiles differed: Chloroflexota were more prominent in hot springs, whereas Campylobacterota made a larger contribution in hydrothermal vents. Acid mine drainage exhibited a different archaeal profile, with notable contributions from Thermoplasmatota and Micrarchaeota. Differences were also evident among habitats assigned to deep isolation. Patescibacteriota were particularly represented in the deep-subsurface collection, whereas Chloroflexota and Nitrospirota contributed more prominently to deep-sea sediment MAGs. Cave MAGs were predominantly bacterial, with substantial contributions from Pseudomonadota, Actinomycetota and Bacillota.

| **Article title** | **Habitat** | **Reference** |
| --- | --- | --- |
| Compendium of 530 metagenome-assembled bacterial and archaeal genomes from the polar Arctic Ocean | Arctic ocean | [1] |
| Metagenome-assembled-genomes recovered from the Arctic drift expedition MOSAiC | Arctic ocean | [2] |
| Metagenomics untangles metabolic adaptations of Antarctic endolithic bacteria at the fringe of habitability | Antarctica saline lake | [3] |
| Unveiling unique microbial nitrogen cycling and nitrification driver in coastal Antarctica | Mixed | [4] |
| A genome and gene catalog of the aquatic microbiomes of the Tibetan Plateau | Mixed | [5] |
| Mapping the metagenomic diversity of the multi-kingdom glacier-fed stream microbiome | Glacier freshwater | [6] |
| Metagenomic insights into microbial community structure and metabolism in alpine permafrost on the Tibetan Plateau | Permafrost | [7] |
| Permafrost microbial communities and functional genes are structured by latitudinal and soil geochemical gradients | Permafrost | [8] |
| Multiple energy sources and metabolic strategies sustain microbial diversity in Antarctic desert soils | Permafrost | [9] |
| A database of glacier prokaryotic genomes and genes for the Three Poles | Glacier | [10] |
| Distinct genes and microbial communities involved in nitrogen cycling between monsoon- and westerlies-dominated Tibetan glaciers | Glacier | [11] |
| Genetic isolation and metabolic complexity of an Antarctic subglacial microbiome | Glacier | [12] |
| Metagenomic insights into novel microbial lineages with distinct ecological functions in the Arctic glacier foreland ecosystems | Glacier | [13] |
| A comprehensive genomic catalog from global cold seeps | Cold seep | [14] |
| Methane filtration and metabolic cooperation of microbial communities in cold seep water columns from South China Sea | Cold seep | [15] |
| Recovery of 1,773 microbial genomes and 2,060 viral genomes from the Mars-analog Qaidam Basin desert | Desert | [16] |
| A deep metagenomic atlas of Qinghai-Xizang Plateau lakes reveals their microbial diversity and salinity adaptation mechanisms | Saline lake | [17] |
| Extremely acidic proteomes and metabolic flexibility in bacteria and highly diversified archaea thriving in geothermal chaotropic brines | Saline lake | [18] |
| Hyperexpansion of genetic diversity and metabolic capacity of extremophilic bacteria and archaea in ancient Andean lake sediments | Saline lake | [19] |
| Mining of novel secondary metabolite biosynthetic gene clusters from acid mine drainage | Acid mine drainage | [20] |
| Patterns and ecological drivers of viral communities in acid mine drainage sediments across Southern China | Acid mine drainage | [21] |
| Covariation of hot spring geochemistry with microbial genomic diversity, function, and evolution | Hot spring | [22] |
| Genome-resolved metagenomic survey of 500 samples from 56 hot springs across the Western US | Hot spring | [23] |
| Recovery of nearly 3,000 archaeal genomes from 152 terrestrial geothermal spring metagenomes | Hot spring | [24] |
| The recovery of 12,789 genomes revealed the diversity, function, and microbial interactions of the geothermal spring microbiome | Hot spring | [25] |
| Expansive microbial metabolic versatility and biodiversity in dynamic Guaymas Basin hydrothermal sediments | Hydrothermal vent | [26] |
| Global deep-sea hydrothermal deposit metagenomes and metagenome-assembled genomes over time and space | Hydrothermal vent | [27] |
| Global patterns of diversity and metabolism of microbial communities in deep-sea hydrothermal vent deposits | Hydrothermal vent | [28] |
| Microbial communities of Auka hydrothermal sediments shed light on vent biogeography and the evolutionary history of thermophily | Hydrothermal vent | [29] |
| A global deep terrestrial biosphere core microbiome | Deep subsurface | [30] |
| Expanded global groundwater microbial diversity reveals bioprospecting potential | Deep subsurface | [31] |
| Metagenome sequencing and 107 microbial genomes from seamount sediments along the Yap and Mariana trenches | Deep sea sediment | [32] |
| Metagenome sequencing and 982 microbial genomes from Kermadec and Diamantina Trenches sediments | Deep sea sediment | [33] |
| Metagenome-assembled genomes of deep-sea sediments: changes in microbial functional potential lag behind redox transitions | Deep sea sediment | [34] |
| Microbiomes in the Challenger Deep slope and bottom-axis sediments | Deep sea sediment | [35] |
| Microbial aerotrophy enables continuous primary production in diverse cave ecosystems | Cave | [36] |
| Reconstruction of 1,979 prokaryotic metagenome-assembled genomes from 37 global cave environments | Cave | [37] |

**Supplementary Table 1 | Habitat classifications of MAG source studies.** Environmental categories were assigned from the source-study descriptions using the supplied classification scheme.

| **Habitat** | **HT** | **LT** | **HS** | **UV** | **LWA** | **EP** | **HM** | **LN** | **HP** | **LO** |
| --- | --- | --- | --- | --- | --- | --- | --- | --- | --- | --- |
| Arctic Ocean |  | ✓ | ✓ | ✓ |  |  |  |  |  |  |
| Antarctica saline lake |  | ✓ | ✓ | ✓ |  | ✓ |  |  |  |  |
| Plateau freshwater |  | ✓ |  | ✓ |  |  |  |  |  |  |
| Glacier freshwater |  | ✓ |  | ✓ |  |  |  | ✓ |  |  |
| Permafrost |  | ✓ |  | ✓ |  | ✓ |  |  |  |  |
| Glacier |  | ✓ |  | ✓ | ✓ |  |  | ✓ |  |  |
| Cold seep |  | ✓ |  |  |  |  |  | ✓ |  | ✓ |
| Desert | ✓ |  |  | ✓ | ✓ | ✓ |  |  |  |  |
| Saline lake |  |  | ✓ |  |  | ✓ |  |  |  |  |
| Acid mine drainage |  |  |  |  |  | ✓ | ✓ |  |  |  |
| Hot spring | ✓ |  |  |  |  | ✓ |  | ✓ |  | ✓ |
| Hydrothermal vent | ✓ |  |  |  |  |  |  | ✓ | ✓ | ✓ |
| Deep subsurface |  | ✓ |  |  |  |  |  | ✓ |  | ✓ |
| Deep-sea sediment |  | ✓ |  |  |  |  |  | ✓ | ✓ | ✓ |
| Cave |  |  |  |  |  |  |  | ✓ |  |  |

**Supplementary Table 2 | Environmental-feature metadata of the 15 isolated habitats.** Binary metadata summarize the dominant environmental constraints assigned to each habitat. Note. HT, high temperature; LT, low temperature; HS, high salinity; UV, high UV radiation; LWA, low water availability; EP, extreme pH; HM, high metal concentrations; LN, low nutrient availability; HP, high pressure; LO, low oxygen availability. A check mark indicates that the environmental feature was assigned to the corresponding habitat according to the predefined classification scheme; a blank cell indicates that the feature was not included in its assigned profile. These symbols are operational habitat-level descriptors rather than direct measurements from every metagenomic sample; therefore, a blank cell does not necessarily indicate the complete environmental absence of a feature.

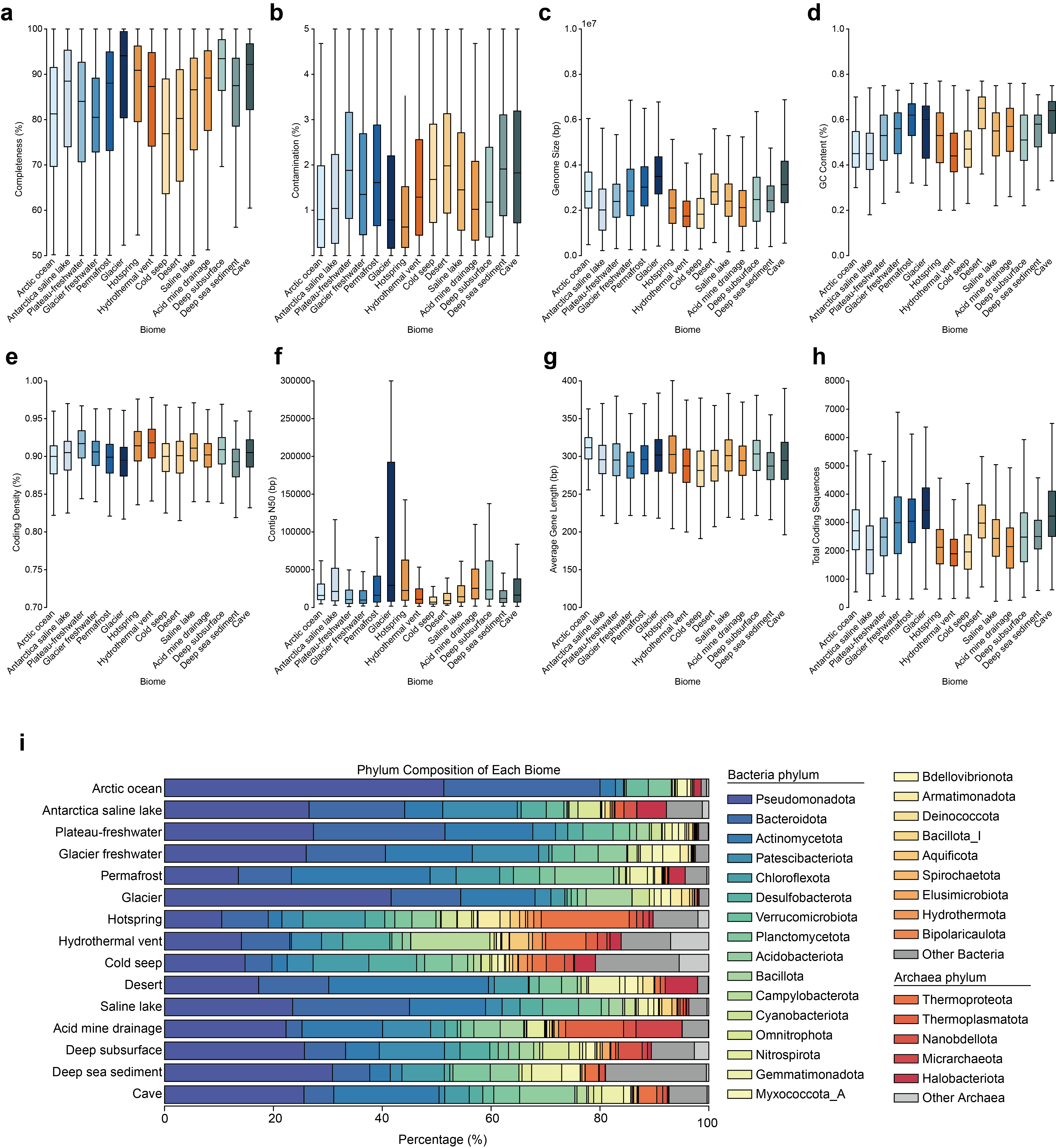

**Supplementary Figure 1 | Quality assessment and taxonomic composition of MAGs recovered from globally distributed isolated microbial habitats. a–h,** Distribution of genome quality metrics for MAGs recovered from different isolated biomes after quality control with CheckM2. Boxplots show the biome-level distributions of **genome completeness** (**a**), **contamination** (**b**), **genome size** (**c**), **GC content** (**d**), **coding density** (**e**), **contig N50** (**f**), **average gene length** (**g**), and **total number of coding sequences** (**h**). Each box represents the interquartile range, the central line indicates the median, and whiskers denote the data range excluding extreme outliers. Biomes include Arctic Ocean, Antarctica saline lake, plateau freshwater, glacier freshwater, permafrost, glacier, hot spring, hydrothermal vent, cold seep, desert, saline lake, acid mine drainage, deep subsurface, deep-sea sediment, and cave environments. These metrics collectively summarize the assembly quality, genome compactness, nucleotide composition, coding potential, and fragmentation level of MAGs across geographically and ecologically diverse isolated habitats. **i,** Relative phylum-level taxonomic composition of MAGs across the 15 isolated biomes. Stacked bars show the percentage of MAGs assigned to each bacterial or archaeal phylum within each biome. Colors indicate phyla, with the remaining bacterial and archaeal assignments grouped as ‘Other Bacteria’ and ‘Other Archaea’, respectively.

### 2. Reference AMP coverage and cross-model predictions

Following smORF prediction and homology screening against reference protein databases, we identified 3,527,632 database-novel smORFs and assessed their similarity to known AMPs and their predictions across eight AMP models. To characterize the relationship between database-novel smORFs and known antimicrobial peptides (AMPs), we constructed an experimentally supported reference dataset by combining 6,169 AMP-positive sequences from the training dataset with additional records from dbAMP3, DRAMP, APD6, AMPDB and DBAASP. All records underwent the same sequence preprocessing and quality-control procedures used for the training dataset. Global sequence-level deduplication yielded 29,420 reference AMPs, with database provenance retained for each sequence.

Searching the 3,527,632 database-novel smORFs against this expanded reference set identified 20,943 sequences with detectable AMP homology (0.59%; Supplementary Fig. 2a). Based on the best MMseqs hit per smORF, sequence identities were concentrated in the 40–60% range, and 64.6% of matched sequences had identities below 60%. Only 721 sequences fell within the 90–100% identity interval. Thus, direct homology to reference AMPs covered only a small fraction of the database-novel sequence space, and most detected matches showed substantial sequence divergence.

Predictions across eight AMP models revealed marked differences in candidate-set sizes and composition (Supplementary Fig. 2b). For example, 621,322 sequences were predicted as AMPs exclusively by amPEPpy and 138,622 exclusively by EvoGradient, whereas 5,986 sequences were jointly predicted by all eight models. These large model-specific subsets and the limited unanimous overlap demonstrate that model choice substantially influences candidate selection within this sequence space.

To examine whether cross-model disagreement varied with habitat breadth, we compared dispersion in model-specific percentile ranks across smORF families with different biome occupancies (Supplementary Fig. 2c). Dispersion varied non-monotonically with biome occupancy. Families restricted to a single biome showed the highest median dispersion, whereas those occurring in an intermediate number of biomes showed lower dispersion. Dispersion increased again among broadly distributed families, although these groups contained substantially fewer families. Together, the limited coverage of reference AMPs and substantial disagreement among prediction models highlight the challenge of prioritizing candidates from database-novel sequence space, particularly among biome-restricted families, and motivate computational prioritization followed by experimental validation.

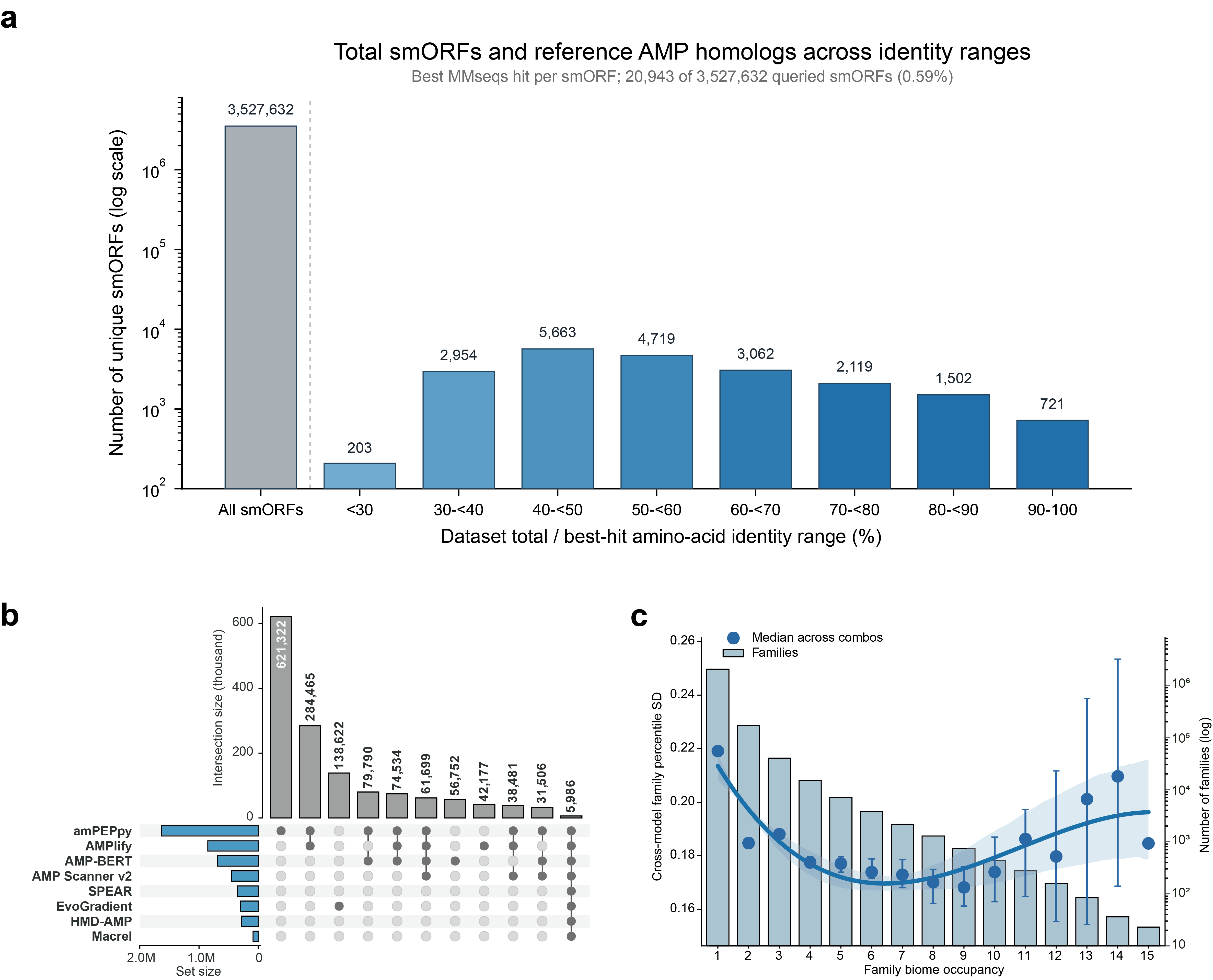

**Supplementary Figure 2 | Sequence identity to reference AMPs, overlap among predictor candidate sets and cross-model prediction disagreement across smORF families a,** Sequence-identity distribution of novel smORFs homologous to reference antimicrobial peptides (AMPs). The grey bar indicates the total number of database-novel smORFs analyzed (3,527,632), whereas the blue bars show the numbers with reported MMseqs hits to reference AMPs across best-hit amino-acid identity ranges. For smORFs with multiple hits, only the highest identity was retained. Exact counts are shown above the bars, and the y axis is displayed on a logarithmic scale. Across the identity bins, 20,943 smORFs (0.59%) had a reference-AMP hit, whereas 3,506,689 (99.41%) had no detected hit under the stated search settings. **b,** Overlap among candidate AMP sets identified by eight predictors: amPEPpy, AMPlify, AMP-BERT, AMP Scanner v2, SPEAR, EvoGradient, HMD-AMP and Macrel. In the UpSet plot, horizontal bars indicate the total candidate-set size for each predictor. Vertical bars show the sizes of the displayed exclusive intersections, with predictor combinations indicated by filled, connected dots below each bar. Numbers above the bars report intersection sizes. **c,** Cross-model prediction disagreement as a function of smORF family habitat breadth. Raw scores from six models (SPEAR, HMD-AMP, AMP-BERT, AMP Scanner, AMPlify and Macrel) were independently converted to percentile ranks across the full analyzed smORF collection to account for differences in score scale. For each family, member-sequence percentile ranks were summarized by their median within each model; the standard deviation of these six model-specific medians defined family-level disagreement.

### 3. Structural diversity of reference antimicrobial peptides and its relationships with sequence physicochemical properties and antimicrobial potency

To characterize structural variation among known antimicrobial peptides (AMPs), we established a contact-map-based framework and examined how the resulting structural groups relate to sequence-derived physicochemical properties and experimentally measured antimicrobial potency. Specifically, we asked how reference AMP structures are distributed across structural groups, which physicochemical properties covary with their structural features, and whether the groups differ in recorded antimicrobial potency. Physicochemical properties and MIC values were not used as clustering inputs.

**Reference structures and contact-map representation.** The reference set comprised 6,169 AMP-positive peptides in the training dataset. Structures were predicted by ESMFold [38]. Accordingly, structural diversity in this analysis refers to the conformations represented in these coordinate models, rather than the complete conformational ensembles of the peptides. Each structure was represented by 31 descriptors derived from its Cα contact map, global geometry and contact graph. A contact was defined between residues separated by more than two sequence positions whose Cα atoms were within 8.0 Å. The descriptors summarized contact abundance, contact sequence separation, spatial compactness and contact-network organization. These descriptors characterize geometric patterns; individual contact-separation features were not interpreted as direct measurements of α-helical or β-strand content.

**Definition and distribution of structural archetypes.** MiniBatch k-means clustering was performed in the transformed and standardized descriptor space for k = 2~30. We selected the number of groups by jointly considering inertia, the Calinski-Harabasz index, the Davies–Bouldin index, mean silhouette coefficient, largest-cluster fraction and structural interpretability (Supplementary Fig. 3a-e). A nine-group solution was selected as a descriptive compromise between within-group compactness, separation and structural resolution. These groups are termed structural archetypes, denoting operational groups in the selected feature space rather than established biological fold classes. UMAP was used only for visualization and did not determine cluster assignments (Supplementary Fig. 3f). Archetypes were numbered in decreasing order of membership, and the observed reference AMP closest to each centroid was selected as its representative (Supplementary Fig. 3g). Membership was uneven: Archetype 1 contained 2,878 AMPs (46.7%), followed by Archetypes 2 and 3 with 1,025 (16.6%) and 855 (13.9%) AMPs, respectively. Together, these three groups comprised 77.1% of the reference set. The remaining six archetypes contained 73~473 AMPs each (1.2~7.7%; Supplementary Table 3).

Inspection of the representative structures, together with the archetype-level contact-map profiles, revealed differences in secondary-structure organization, chain extension and non-local packing (Supplementary Figs. 3g and 4a). Archetypes 1, 3 and 4 displayed distinct helix-containing conformations. The representative of Archetype 1 contained a local helical segment accompanied by extended and loop regions, consistent with a contact profile dominated by local rather than long-range interactions. Archetype 3 combined a helical segment with a long-extended region and showed relatively large radii of gyration and terminal distances. By comparison, Archetype 4 was predominantly helical, with pronounced enrichment of near-diagonal contacts. Archetypes 2, 7, 8 and 9 exhibited different forms of β-strand-containing organization. Archetype 2 was represented by a compact mixed α/β conformation, whereas Archetype 7 contained several β-strand segments connected by loops. Both groups showed relatively high mean contact degrees and contact-graph clustering, consistent with more extensively interconnected contact networks. Archetype 8 displayed a more elongated, β-rich conformation with a less densely connected contact network than Archetypes 2 and 7. Archetype 9 was represented by a β-hairpin-like conformation comprising two adjacent antiparallel strands, accompanied by prominent off-diagonal contacts and a high terminal-contact fraction. The remaining archetypes were characterized by limited packing. Archetype 5 showed an extended, contact-poor conformation, with low contact density, large length-normalized geometric dimensions and a highly disconnected contact graph. Archetype 6 was represented by a predominantly loop-like conformation with limited local secondary structure and comparatively sparse contacts. These descriptions refer to the analyzed coordinate models; extended or sparsely connected conformations were not interpreted as evidence of intrinsic disorder.

Together, these profiles indicate that the AMP-positive training set spans variation in local secondary-structure organization and non-local packing, including predominantly helical, mixed α/β, β-rich and extended conformations. The archetype frequencies describe this training-set collection, whereas the representative structures illustrate characteristic conformations without implying structural uniformity within each group.

**Relationships with sequence physicochemical properties.** We next examined how the structural archetypes were associated with sequence-derived physicochemical properties. Archetype-level profiles revealed differences in hydrophobicity-related indices, charge-related properties and sequence length (Supplementary Fig. 4b). The helix-containing Archetypes 1 and 4 showed relatively high GRAVY values and hydrophobic moments, together with low Boman and instability indices. Archetype 1 additionally exhibited relatively high isoelectric point, charge and aliphatic index, whereas Archetype 4 showed a particularly high GRAVY value. By contrast, the elongated, partially helical Archetype 3 displayed lower GRAVY, hydrophobicity and hydrophobic moment, alongside higher Boman and instability indices. Thus, the presence of a helical segment did not correspond to a uniform physicochemical profile across archetypes.

The compact Archetypes 2 and 7, characterized by extensive non-local contacts, contained relatively long peptides with lower isoelectric points, charge, aromaticity and aliphatic indices. However, these properties were not shared by all β-strand-containing archetypes. The β-hairpin-like Archetype 9 showed relatively high aromaticity and charge, distinguishing it from Archetypes 2 and 7. Conversely, the extended, contact-poor Archetype 5 also exhibited relatively high isoelectric point and charge, together with a high instability index. These comparisons indicate that similar charge-related properties can accompany different conformations, while distinct β-strand-containing architectures can have different sequence physicochemical profiles. Spearman correlation analysis across individual AMPs further identified relationships between physicochemical properties and contact organization (Supplementary Fig. 4c). GRAVY, aliphatic index and isoelectric point were positively associated with short-range contact fractions and negatively associated with medium- and long-range contact fractions and relative contact order. Hydrophobic moment was positively associated with near-diagonal contact density and enrichment, as well as overall contact density and mean contact degree. These patterns linked several hydrophobicity- and charge-related sequence properties to local contact organization, although the different hydrophobicity-related indices did not show identical correlation profiles. Higher Boman indices were associated with lower contact density and larger radii of gyration, whereas instability indices were associated with lower contact density and a higher connected-component ratio.

Sequence length showed a distinct correlation pattern: longer peptides had higher mean contact degrees, greater contact sequence separations, larger radii of gyration and more extensive contact-graph connectivity, but lower overall contact densities and short-range contact fractions. Length therefore represents an important covariate when interpreting the observed structural-physicochemical relationships. Together, these analyses identify physicochemical differences among the structural archetypes and continuous associations across individual AMPs, while indicating that no single sequence property uniquely specifies an archetype.

**Relationships with peptide antimicrobial potency.** To examine whether structural archetypes corresponded to differences in antimicrobial potency, we compared MIC records from GRAMPA across the nine archetypes, separately for *Staphylococcus aureus* and *Escherichia coli* (Supplementary Fig. 4d). Potency was displayed as −log₂(MIC), with larger values indicating lower MICs and greater potency. For both bacterial species, MIC-derived potency distributions overlapped substantially across archetypes, with considerable variation within individual groups. The extended, contact-poor Archetype 5 overlapped in recorded potency with the helix-containing Archetypes 1 and 4 and the more compact Archetypes 2 and 7. Thus, greater structural compactness or the presence of a particular secondary-structure arrangement did not correspond to a consistent potency advantage. Some archetype- and species-dependent shifts were nevertheless apparent. For example, the β-hairpin-like Archetype 9 tended to show lower potency against E. coli, although its distribution still overlapped those of other archetypes. Together, these observations suggest that the structural archetypes capture differences in peptide organization without clearly separating high- and low-potency AMPs. Broad structural category alone therefore provides limited resolution for distinguishing antimicrobial potency within this AMP collection.

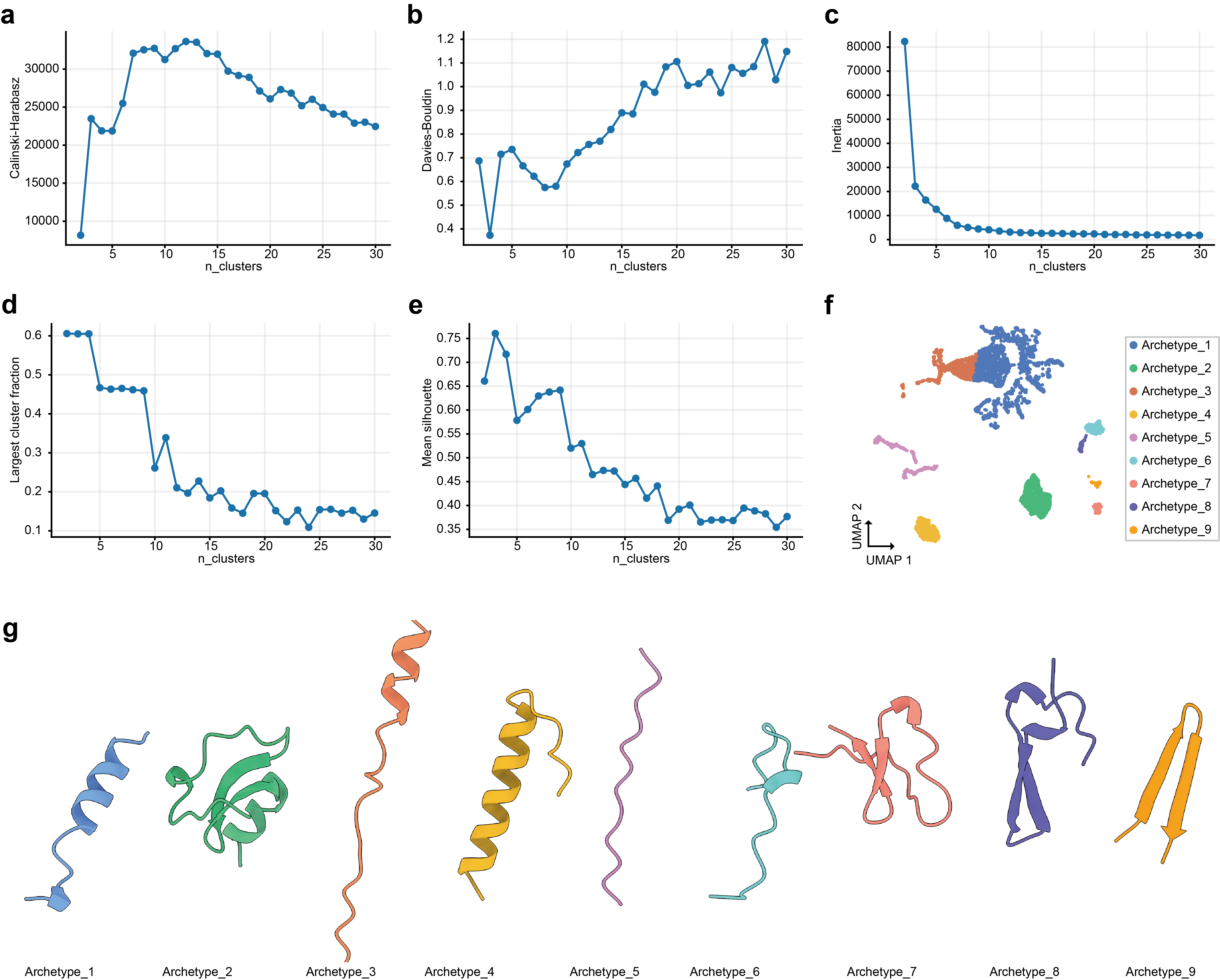

**Supplementary Figure 3 | Contact-map-based clustering resolves nine AMP structural archetypes. a–e**, Parameter search for the number of AMP structural clusters based on tertiary-structure contact-map features. Cluster quality was evaluated using the Calinski–Harabasz index **(a),** Davies–Bouldin index **(b),** inertia **(c),** largest cluster fraction **(d)**, and mean silhouette score **(e).** These metrics jointly assess cluster separation, compactness, within-cluster variance, cluster-size imbalance, and overall clustering coherence. Based on their combined behavior, n = 9 was selected as a balanced solution that captures structural diversity without excessive fragmentation. **f,** UMAP visualization of AMP structures using contact-map-derived features, colored by the nine inferred structural archetypes. **g,** Representative tertiary structures of the nine AMP archetypes, showing distinct helical, extended, β-strand-containing, and mixed structural topologies.

| **Archetype** | **Cluster size** | **Mean distance to centroid** | **Distance Q50** | **Distance Q75** | **Distance Q90** | **Distance Q95** | **Threshold quantile** | **Distance threshold** |
| --- | --- | --- | --- | --- | --- | --- | --- | --- |
| Archetype_1 | 2,878 | 0.605 | 0.547 | 0.775 | 1.048 | 1.191 | 0.95 | 1.191 |
| Archetype_2 | 1,025 | 0.786 | 0.673 | 0.946 | 1.311 | 1.617 | 0.95 | 1.617 |
| Archetype_3 | 855 | 0.879 | 0.860 | 1.066 | 1.316 | 1.584 | 0.95 | 1.584 |
| Archetype_4 | 473 | 0.902 | 0.850 | 1.073 | 1.359 | 1.586 | 0.95 | 1.586 |
| Archetype_5 | 425 | 0.155 | 0.125 | 0.194 | 0.258 | 0.366 | 0.95 | 0.366 |
| Archetype_6 | 214 | 1.144 | 1.121 | 1.392 | 1.667 | 1.784 | 0.95 | 1.784 |
| Archetype_7 | 126 | 0.679 | 0.517 | 0.725 | 1.224 | 1.417 | 0.95 | 1.417 |
| Archetype_8 | 100 | 1.285 | 1.112 | 1.347 | 1.978 | 3.091 | 0.95 | 3.091 |
| Archetype_9 | 73 | 1.621 | 0.703 | 2.681 | 3.268 | 3.955 | 0.95 | 3.955 |

**Supplementary Table 3 | Summary statistics for AMP structural archetypes.** Centroid-distance distributions and cluster-specific thresholds for the nine AMP structural archetypes. Distance threshold is defined by the listed threshold quantile of the within-archetype centroid-distance distribution.

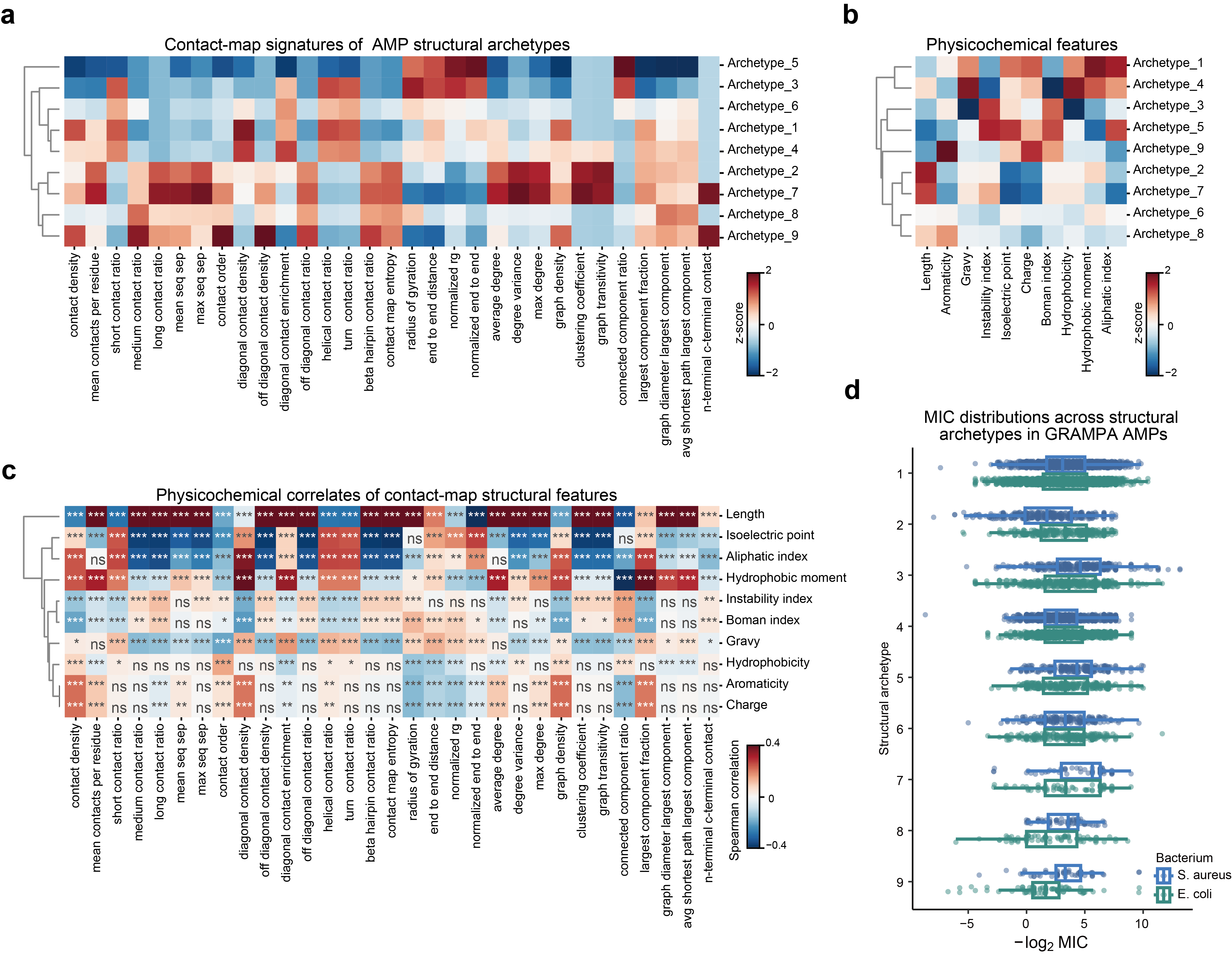

**Supplementary Figure 4 | Contact-map-defined AMP structural archetypes are associated with physicochemical properties.** Antimicrobial peptides (AMPs) were assigned to nine structural archetypes using descriptors derived from tertiary-structure contact maps. **a,** Heat map showing the standardized contact-map descriptor profiles of the nine archetypes, encompassing contact density and sequence separation, contact-order measures, and graph-topological properties. **b,** Standardized physicochemical profiles of the same archetypes. In a and b, colors represent z scores (red, higher; blue, lower), and dendrograms indicate hierarchical clustering of the archetypes. **c,** Spearman rank correlations between contact-map descriptors and physicochemical properties across the AMPs. Colors represent Spearman’s ρ, with red and blue indicating positive and negative correlations, respectively. *P < 0.05, **P < 0.01 and ***P < 0.001; ns, not significant. **d,** Distributions of −log₂-transformed minimum inhibitory concentration (MIC) values recorded in GRAMPA for *Staphylococcus aureus* and *Escherichia coli*, stratified by structural archetype. Each point represents one GRAMPA MIC record [39].

### 4. Peptide representation evaluation

To evaluate how different peptide representations capture antimicrobial activity labels and structural organization, we compared one-hot sequence encoding, 1–2-mer composition, physicochemical features (PhysChem), ESM2-650M embeddings, SaProt embeddings and the fused representation used by SPEAR. We assessed AMP/non-AMP discrimination across 13,652 peptides and structural-archetype discrimination within the AMP subset, using classification performance, pairwise separation, nearest-neighbor enrichment and clustering metrics (Supplementary Fig. 5 and Supplementary Tables 4-5).

**Representation of AMP/non-AMP labels.** Protein language model embeddings achieved higher cross-validated classification performance than the conventional representations. The fused SPEAR representation yielded the highest mean AUROC (0.922 ± 0.006), AUPR (0.928 ± 0.005), balanced accuracy (0.855 ± 0.006) and F1 score (0.838 ± 0.008), although its numerical improvements over ESM2-650M alone were small (Supplementary Table 4). Local neighborhood analyses showed a different ordering: ESM2-650M consistently exhibited the strongest AMP/non-AMP enrichment, followed by SPEAR and SaProt. For example, macro-adjusted enrichment at k = 5 was 0.596 for ESM2-650M, 0.582 for SPEAR and 0.562 for SaProt, compared with 0.370–0.518 for the conventional representations. Similar trends were observed for macro- and micro-averaged same-label neighbor fractions (Supplementary Fig. 5a–d). Thus, fusion retained strong local organization of AMP/non-AMP labels without improving neighborhood homogeneity beyond ESM2-650M alone.

**Representation of AMP structural archetypes.** SaProt showed the strongest structural-archetype organization, with a pairwise AUROC of 0.827 (95% CI, 0.821–0.833), compared with 0.668 (0.661–0.675) for ESM2-650M. Fusion increased this value to 0.769 (0.762–0.775), indicating greater structural-archetype separation than ESM2-650M alone, while remaining below SaProt (Supplementary Table 5). This ordering was consistent across local enrichment and archetype-classification metrics. Macro-adjusted enrichment at k = 5 was 0.527 for SaProt, 0.487 for SPEAR and 0.368 for ESM2-650M; corresponding nearest-neighbor balanced accuracies were 0.593, 0.548 and 0.444. SaProt also achieved the highest Calinski–Harabasz index and lowest Davies–Bouldin index. However, silhouette scores remained close to zero, including for SaProt, indicating substantial overlap among structural archetypes rather than uniformly discrete clusters.

Across both analyses, adjusted enrichment and same-label or same-archetype neighbor fractions decreased as neighborhood size increased, showing that label-associated organization was strongest at local scales. UMAP projections provided a qualitative view of archetype-associated regions and their overlap. Together, these results support complementary information content in ESM2-650M and SaProt embeddings: ESM2-650M more strongly organized AMP/non-AMP labels, whereas SaProt more strongly organized structural archetypes. Their fusion retained strong AMP-label organization while increasing structural-archetype organization relative to ESM2-650M alone, providing a representation that captures both aspects of peptide variation.

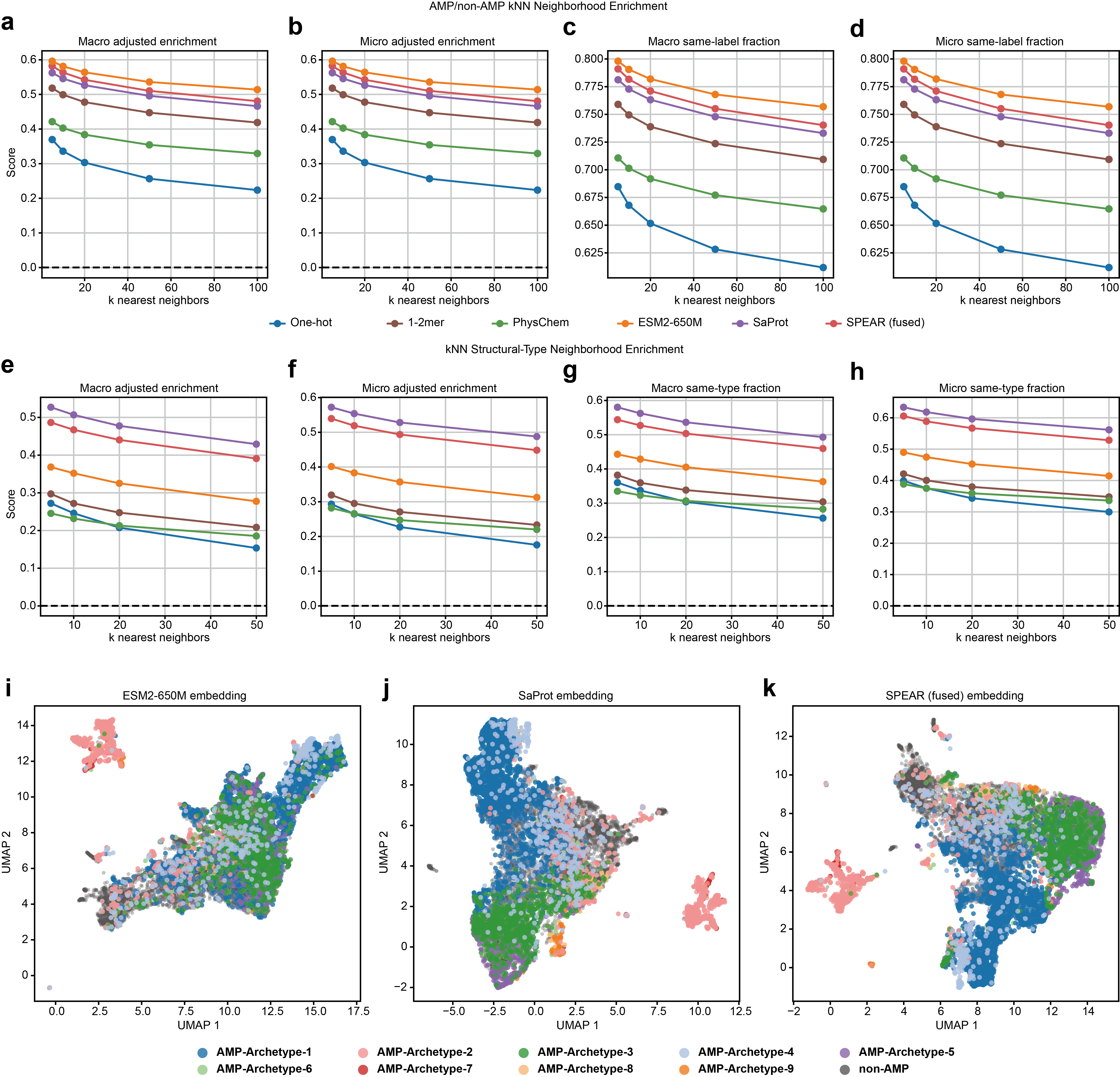

**Supplementary Figure 5 | SPEAR retains complementary AMP-label and structural-archetype information in its fused embedding.** Antimicrobial peptides (AMPs) and non-AMP sequences were represented using one-hot sequence encoding, 1–2-mer composition, physicochemical features (PhysChem), ESM2-650M embeddings, SaProt embeddings or fused SPEAR embeddings. For each query peptide, local organization in the corresponding representation space was evaluated across increasing numbers of nearest neighbors, *k*. **a–d,** Macro- and micro-averaged adjusted enrichment (**a, b**) and same-label neighbor fractions (**c, d**) for AMP/non-AMP labels. **e–h,** Corresponding macro- and micro-averaged adjusted enrichment (**e, f**) and same-archetype neighbor fractions (**g, h**) for the nine contact-map-defined AMP structural archetypes. Macro averages assign equal weight to each label or archetype, whereas micro averages pool all query peptides. Higher scores indicate stronger local neighborhood homogeneity, and dashed horizontal lines denote zero adjusted enrichment. ESM2-650M yielded the highest observed AMP/non-AMP neighborhood-enrichment scores, whereas SaProt yielded the highest structural-archetype enrichment. SPEAR remained among the strongest representations in both analyses, consistent with the retention of complementary sequence- and structure-associated information after fusion. Enrichment decreased with increasing *k* for all representations, indicating that label and archetype organization was strongest at local neighborhood scales. **i-k,** Two-dimensional uniform manifold approximation and projection (UMAP) visualizations of the ESM2-650M (**i**), SaProt (**j**) and fused SPEAR (**k**) embeddings. Each point represents one peptide, coloured according to its AMP structural archetype; non-AMP sequences are shown in grey.

| Metric | One-hot | 1-2-mer | PhysChem | ESM2-650M | SaProt | SPEAR |
| --- | --- | --- | --- | --- | --- | --- |
| CV AUROC ↑ | 0.861 ± 0.008 | 0.878 ± 0.009 | 0.826 ± 0.008 | **0.920 ± 0.006** | 0.915 ± 0.006 | **0.922 ± 0.006** |
| CV AUPR ↑ | 0.867 ± 0.008 | 0.841 ± 0.011 | 0.799 ± 0.009 | **0.926 ± 0.006** | 0.923 ± 0.005 | **0.928 ± 0.005** |
| CV balanced accuracy ↑ | 0.792 ± 0.008 | 0.811 ± 0.010 | 0.772 ± 0.009 | **0.853 ± 0.007** | 0.849 ± 0.007 | **0.855 ± 0.006** |
| CV F1 ↑ | 0.771 ± 0.009 | 0.795 ± 0.011 | 0.738 ± 0.012 | **0.836 ± 0.009** | 0.831 ± 0.008 | **0.838 ± 0.008** |
| CV MCC ↑ | 0.586 | 0.620 | 0.555 | **0.714** | 0.706 | **0.718** |
| Pairwise AUROC (95% CI) ↑ | 0.526 (0.520-0.533) | 0.555 (0.550-0.559) | 0.542 (0.538-0.549) | **0.564 (0.560-0.569)** | 0.550 (0.545-0.555) | **0.555 (0.549-0.560)** |
| Pairwise AUPR (95% CI) ↑ | 0.530 (0.522-0.537) | 0.557 (0.551-0.563) | 0.542 (0.536-0.549) | **0.577 (0.570-0.584)** | 0.559 (0.552-0.566) | **0.564 (0.555-0.571)** |
| Rank-biserial effect ↑ | 0.052 | 0.110 | 0.085 | **0.129** | 0.101 | **0.110** |
| Delta median ↑ | 0.003 | 0.026 | **0.221** | **0.089** | 0.047 | 0.046 |
| AE@5 ↑ | 0.370 | 0.518 | 0.421 | **0.596** | 0.562 | **0.582** |
| AE@10 ↑ | 0.336 | 0.499 | 0.403 | **0.581** | 0.546 | **0.563** |
| AE@20 ↑ | 0.303 | 0.478 | 0.384 | **0.564** | 0.526 | **0.542** |
| AE@50 ↑ | 0.257 | 0.447 | 0.354 | **0.536** | 0.496 | **0.510** |
| AE@100 ↑ | 0.224 | 0.419 | 0.329 | **0.514** | 0.466 | **0.481** |
| Silhouette ↑ | 0.005 | 0.044 | 0.058 | **0.074** | 0.053 | **0.055** |
| Calinski-Harabasz index ↑ | 35.0 | 283.6 | **631.6** | **542.3** | 389.2 | 399.2 |
| Davies-Bouldin index ↓ | 19.647 | 6.884 | **4.373** | **4.983** | 5.889 | 5.816 |

**Supplementary Table 4 | Benchmark of AMP and non-AMP representation methods** All representations used 13652 samples and 40000 pairwise comparisons. Cosine distance was used for all representations except Physchem, which used Euclidean distance. AE, macro-adjusted enrichment; CV, cross-validation; MCC, Matthews correlation coefficient.

| Metric | One-hot | 1-2-mer | PhysChem | ESM2-650M | SaProt | SPEAR |
| --- | --- | --- | --- | --- | --- | --- |
| AUROC (95% CI) ↑ | 0.564 (0.557-0.573) | 0.609 (0.601-0.616) | 0.611 (0.604-0.619) | 0.668 (0.661-0.675) | **0.827 (0.821-0.833)** | **0.769 (0.762-0.775)** |
| AUPR ↑ | 0.573 | 0.635 | 0.615 | 0.669 | **0.818** | **0.768** |
| Rank-biserial effect ↑ | 0.127 | 0.217 | 0.223 | 0.335 | **0.654** | **0.537** |
| Delta median ↑ | 0.010 | 0.062 | **0.601** | 0.233 | **0.420** | 0.313 |
| AE@5 ↑ | 0.272 | 0.297 | 0.245 | 0.368 | **0.527** | **0.487** |
| AE@10 ↑ | 0.246 | 0.272 | 0.232 | 0.352 | **0.507** | **0.467** |
| AE@20 ↑ | 0.208 | 0.247 | 0.213 | 0.325 | **0.478** | **0.440** |
| AE@50 ↑ | 0.154 | 0.208 | 0.185 | 0.277 | **0.429** | **0.391** |
| BA KNN ↑ | 0.362 | 0.378 | 0.329 | 0.444 | **0.593** | **0.548** |
| F1 KNN ↑ | 0.388 | 0.390 | 0.341 | 0.465 | **0.607** | **0.562** |
| BA NC ↑ | 0.435 | 0.420 | 0.338 | 0.438 | **0.614** | **0.585** |
| F1 NC ↑ | 0.397 | 0.341 | 0.306 | 0.321 | **0.522** | **0.483** |
| Micro silhouette ↑ | -0.000 | -0.037 | -0.062 | -0.088 | **0.028** | -0.015 |
| Macro silhouette ↑ | 0.001 | -0.037 | -0.098 | -0.051 | **0.022** | -0.002 |
| Balanced silhouette ↑ | -0.000 | -0.039 | -0.064 | -0.057 | **0.016** | -0.007 |
| Calinski-Harabasz index ↑ | 11.4 | 61.2 | 165.0 | 132.3 | **299.3** | **206.4** |
| Davies-Bouldin index ↓ | 15.163 | 9.381 | 7.029 | 7.670 | **4.470** | **5.578** |
| Negative silhouette clusters ↓ | 62.5% | 87.5% | 75.0% | 75.0% | **37.5%** | **62.5%** |

**Supplementary Table 5 | Benchmark of AMP structure representations**. All representations used 6096 samples and 8 clusters after excluding Archetype 9 (n = 73) (preprocessing: auto; minimum cluster size: 100; silhouette sample size: 5000; maximum per cluster: 500). Cosine distance was used for all representations except PhysChem, which used Euclidean distance. AE, macro-adjusted enrichment; BA, balanced accuracy; NC, nearest centroid.

### 5. Model ablation study

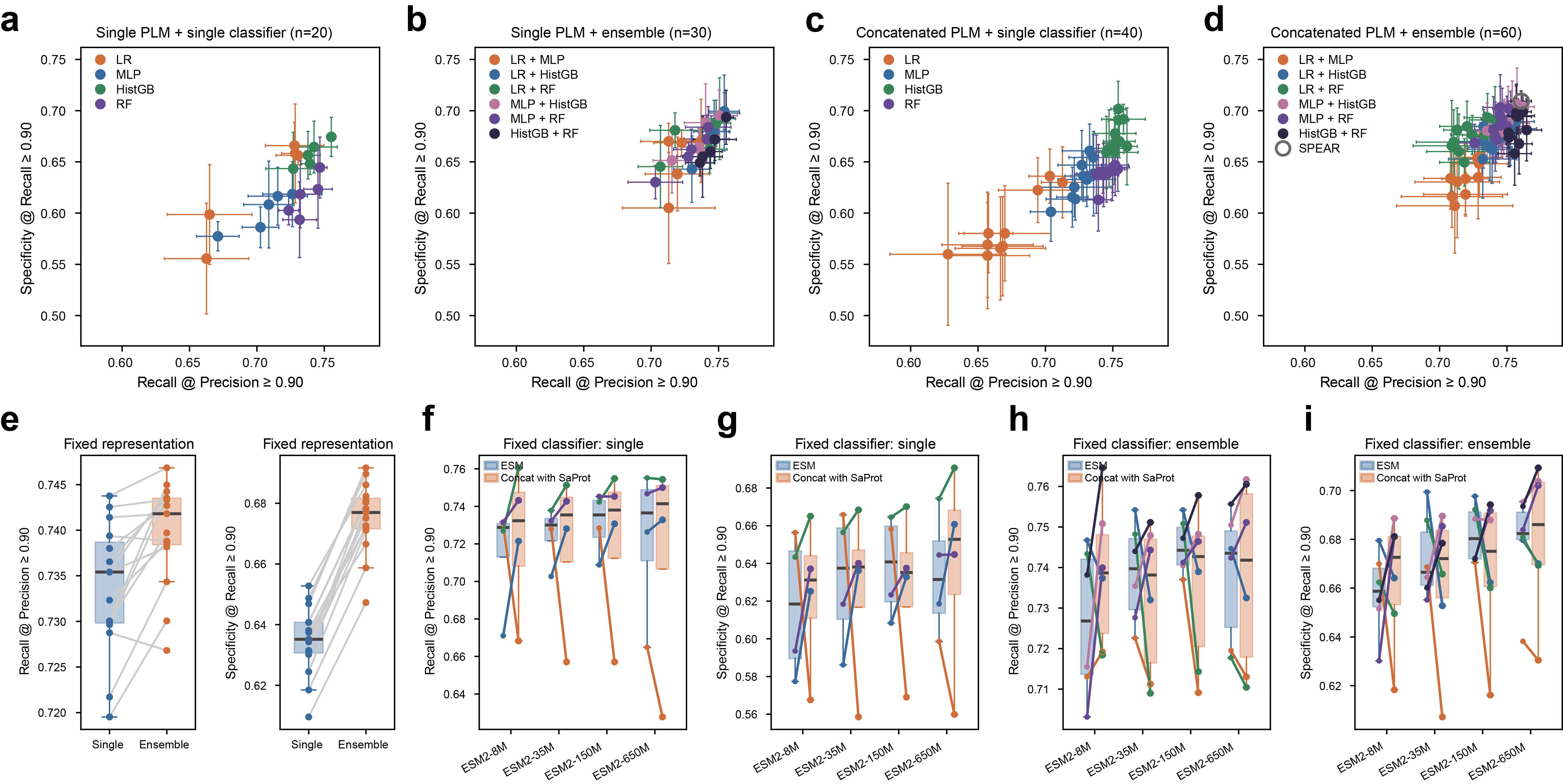
**Supplementary Figure 6 | Ablation of protein language-model representations and classifiers under a 30% sequence-identity constraint.** All configurations were evaluated using the same five held-out test folds. For each fold, training sequences sharing ≥30% sequence identity with any held-out test sequence were removed, while the test set remained unchanged. **a–d,** Performance of individual representation–classifier configurations combining a single protein language model (PLM) with a single classifier (**a**, 20 configurations), a single PLM with a pairwise classifier ensemble (**b**, 30 configurations), a concatenated PLM pair with a single classifier (**c**, 40 configurations), or a concatenated PLM pair with a pairwise classifier ensemble (**d**, 60 configurations). The five PLMs were ESM2-8M, ESM2-35M, ESM2-150M, ESM2-650M and SaProt. The x-axis shows the five-fold mean of the maximum recall achieved at precision ≥0.90; the y-axis shows the five-fold mean of the maximum specificity achieved at recall ≥0.90. Error bars indicate standard deviations across the five folds, and colors denote classifiers or classifier ensembles. The open circle in **d** denotes SPEAR, which concatenates ESM2-650M and SaProt embeddings and averages the positive-class probabilities predicted by HistGradientBoosting and Random Forest. All four panels share the same axis limits. **e,** Comparison of single classifiers and pairwise classifier ensembles with representation held fixed. Each connected pair represents one of the 15 representations comprising five individual PLM embeddings and ten concatenated PLM pairs. Values are the median, across the four single classifiers or six pairwise ensembles, of the five-fold mean recall at precision ≥0.90 (left) or specificity at recall ≥0.90 (right). Boxes summarize these representation-level medians. **f–i,** Effect of concatenating SaProt embeddings with each ESM2 embedding while holding the classifier configuration fixed. Panels show recall at precision ≥0.90 (**f, h**) and specificity at recall ≥0.90 (**g, i**) for single classifiers (**f, g**) and pairwise classifier ensembles (**h, i**). Each line connects results for the same classifier configuration using ESM2 alone (filled point) or ESM2 concatenated with SaProt (open point). Blue and orange boxes summarize classifier-level results for the two representation conditions, respectively. LR, logistic regression; MLP, multilayer perceptron; HistGB, histogram-based gradient boosting; RF, random forest.

### 6. Validation and optimization of liquid distribution in the MicroGrad microfluidic chip

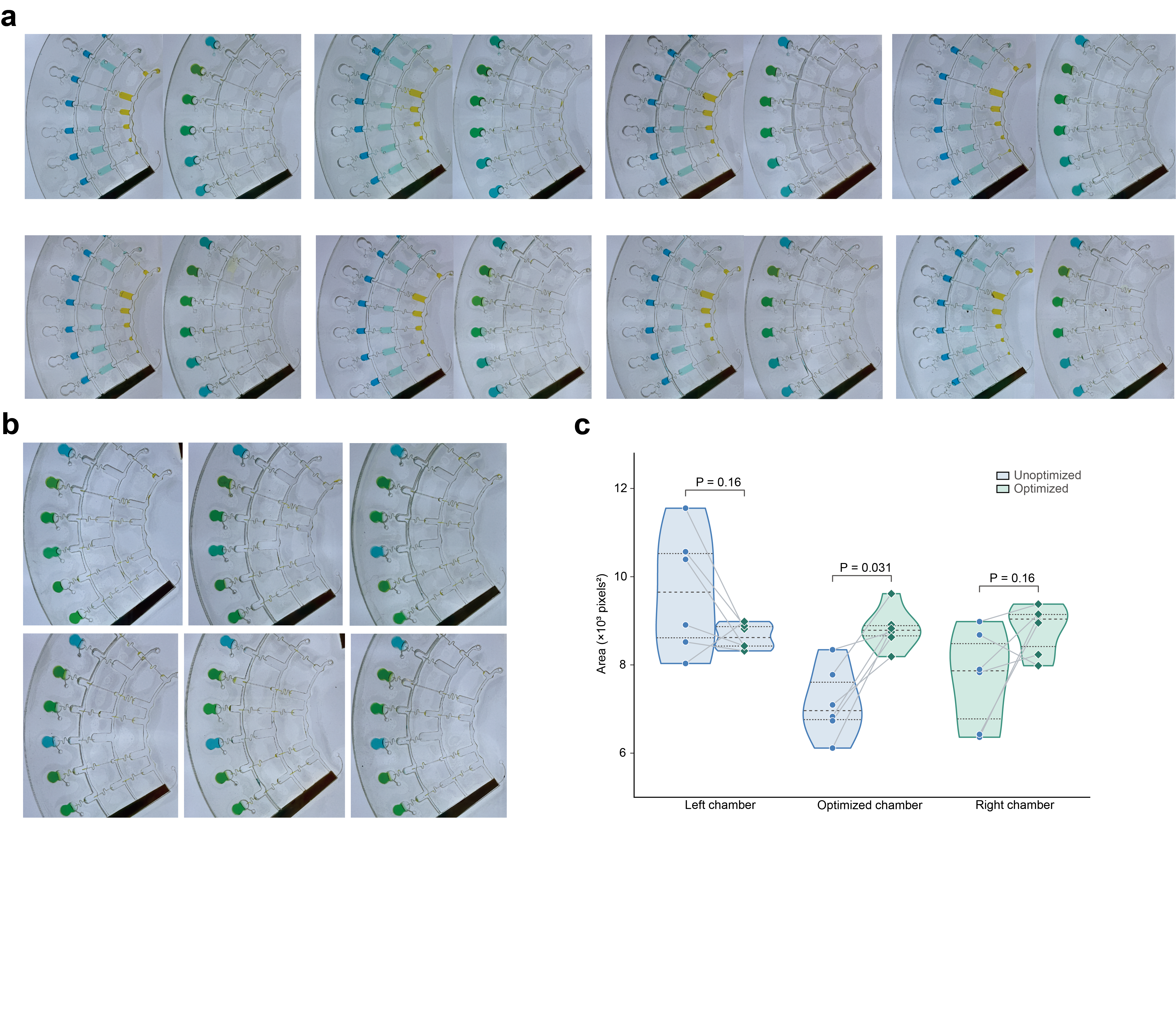

**Supplementary Fig. 7 | Dye-based characterization of liquid distribution and hydraulic resistance optimization in the MicroGrad chip.** **a,** Images of eight chips loaded with dye solutions before (left in each pair) and after (right) centrifugation. **b,** Representative images showing dye distribution in chips with unoptimized and optimized hydraulic resistance. **c,** Quantification of the dye-filled area in the left chamber, the chamber targeted for optimization and the right chamber for the unoptimized and optimized designs. Points represent individual measurements, and lines connect paired observations. Violin plots show the distributions of the measurements. P values were calculated using two-sided paired Wilcoxon signed-rank tests.

### 7. Properties of experimentally validated peptides

To characterize the secondary structures of the validated AMPs, we compared their circular dichroism (CD) spectra in water and a membrane-mimetic environment containing 10 mM SDS. The peptides predominantly exhibited disordered and antiparallel β-sheet-like signatures in water, whereas SDS generally increased their α-helical content (Fig. 4h, Supplementary Fig. 8a). However, the increase in helicity (Δα = α[SDS] − α[water]) was not significantly associated with MIC-derived antimicrobial potency (Supplementary Fig. 8b-i), indicating that the extent of helix induction alone did not explain differences in activity.

We further examined peptide self-assembly in PBS using dynamic light scattering (DLS) and atomic force microscopy (AFM). DLS revealed concentration-dependent formation of nanoscale assemblies (Supplementary Fig. 9a), while AFM resolved diverse morphologies, including nanofibres, nanospheres and hook-like structures, with heights of 8~30 nm and apparent lateral widths of 80~150 nm (Supplementary Fig. 9b-e). Together, these measurements support peptide self-assembly in PBS, although its contribution to the β-sheet-like CD signatures observed in water remains to be established.

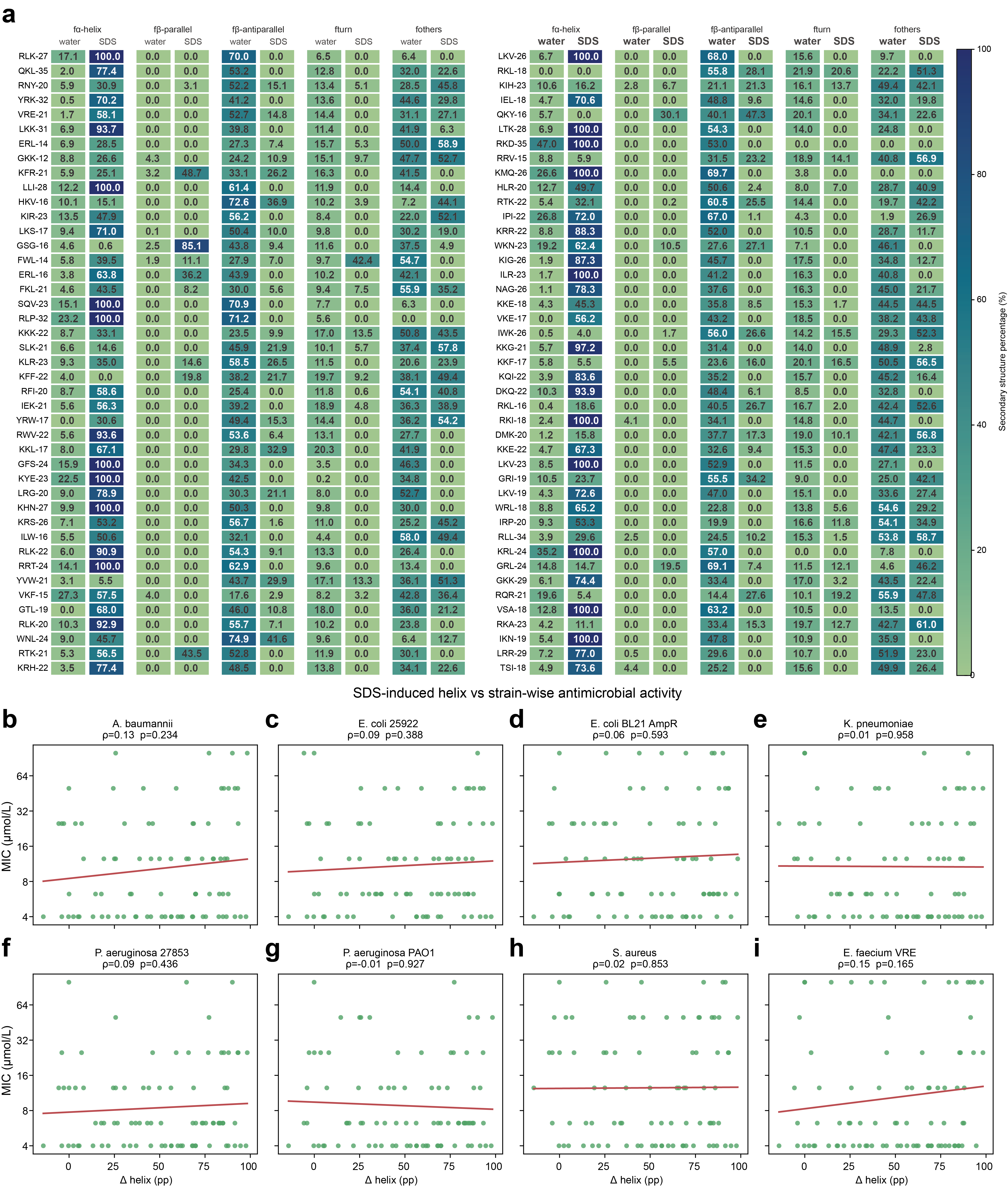

**Supplementary Figure 8 | Secondary-structure compositions of all synthesized peptides in water and sodium dodecyl sulfate (SDS) and their associations with strain-specific antimicrobial activity. a,** Percentages of α-helix, parallel β-sheet, antiparallel β-sheet, turn and other structures in water and SDS, estimated by deconvolution of circular dichroism spectra using the BeStSel web server. **b–i,** Relationships between the SDS-induced change in α-helix content (Δ helix, percentage points (pp)) and the strain-specific minimum inhibitory concentration (MIC) for *A. baumannii* (**b**), *E. coli* 25922 (**c**), *E. coli* BL21 AmpR (**d**), *K. pneumoniae* (**e**), *P. aeruginosa* 27853 (**f**), *P. aeruginosa* PAO1 (**g**), *S. aureus* (**h**) and vancomycin-resistant *E. faecium* (VRE; **i**). Panels **b–g** represent Gram-negative bacteria, whereas panels **h,i** represent Gram-positive bacteria. Each point represents one synthesized peptide, and the red lines indicate fitted trends. Spearman’s rank correlation coefficients (ρ) and corresponding P values are shown in each panel. None of the correlations was statistically significant (all P > 0.05).

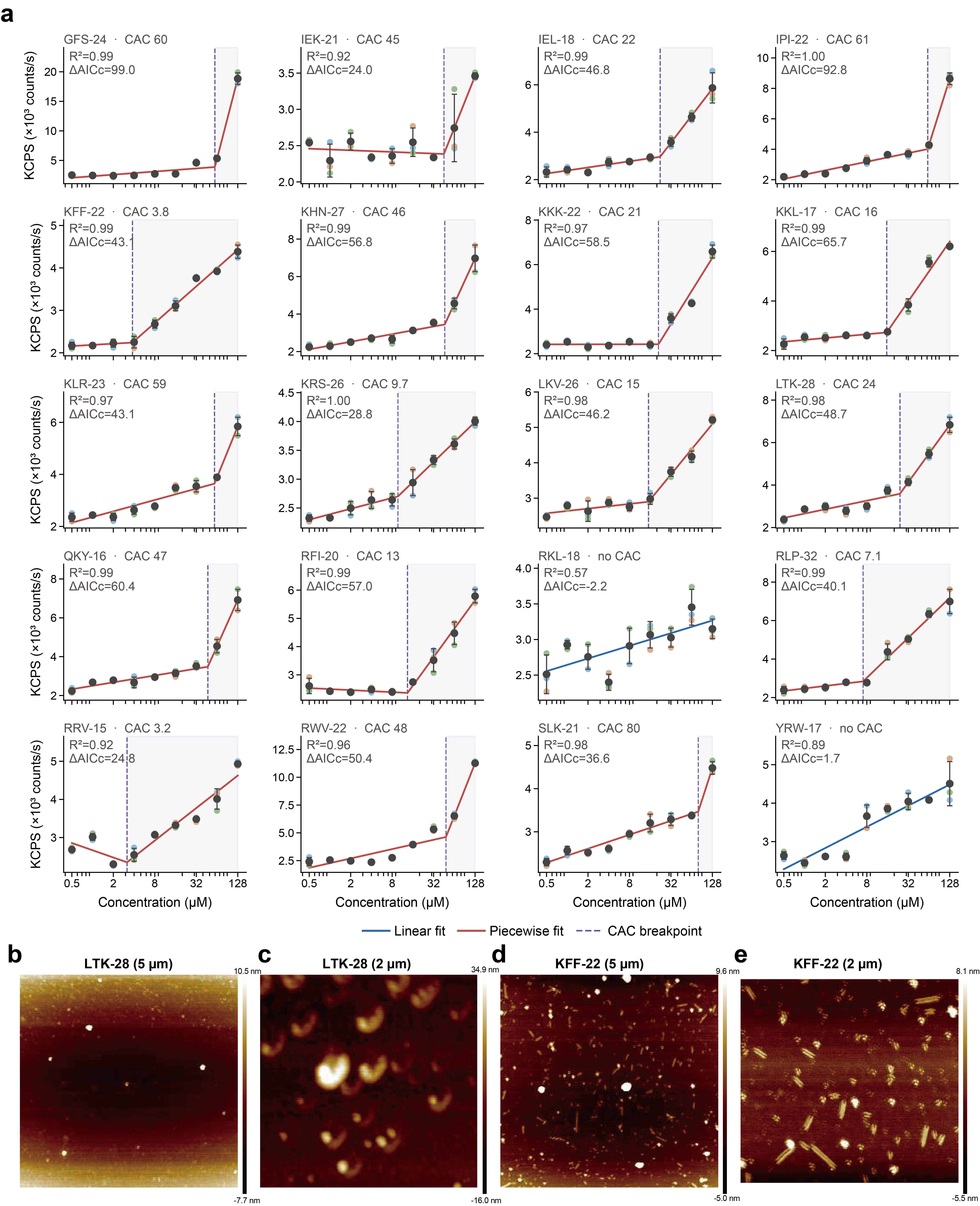

**Supplementary Figure 9 | Concentration-dependent aggregation and self-assembly of selected AMPs. a,** Dynamic light scattering count rates across peptide concentrations. Red and blue lines indicate piecewise and linear fits, respectively; dashed purple lines mark estimated critical aggregation concentrations (CACs). **b-e,** Atomic force microscopy height images of LTK-28 and KFF-22 over 5-μm and 2-μm scan fields, showing diverse nanoscale assemblies. Colour scales indicate height.

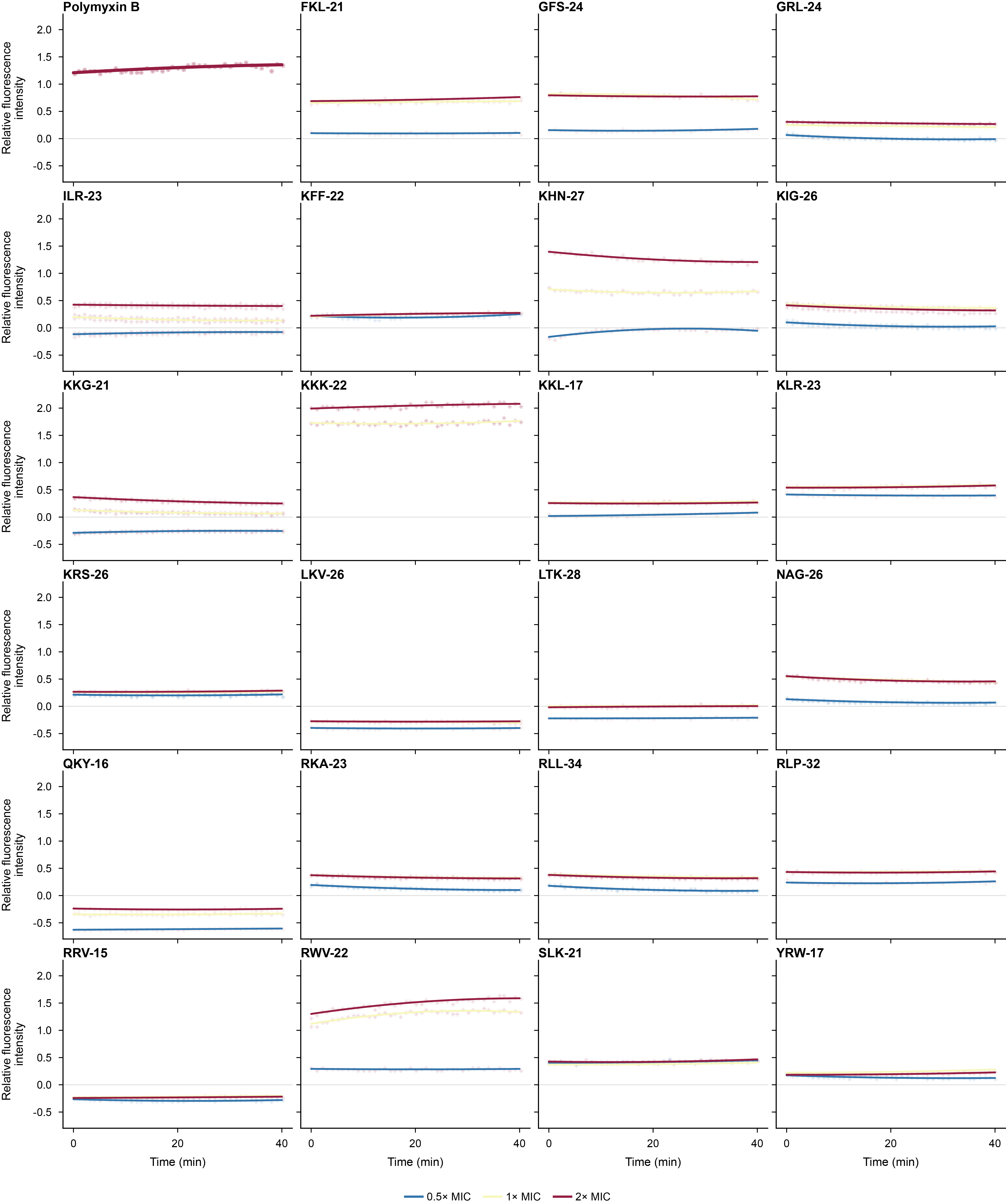

**Supplementary Figure 10 | Outer-membrane permeabilization of *Escherichia coli* measured using the N-phenyl-1-naphthylamine (NPN) uptake assay.** Relative fluorescence was monitored for 40 min after treatment with the indicated peptides at 0.5× MIC, 1× MIC, and 2× MIC.

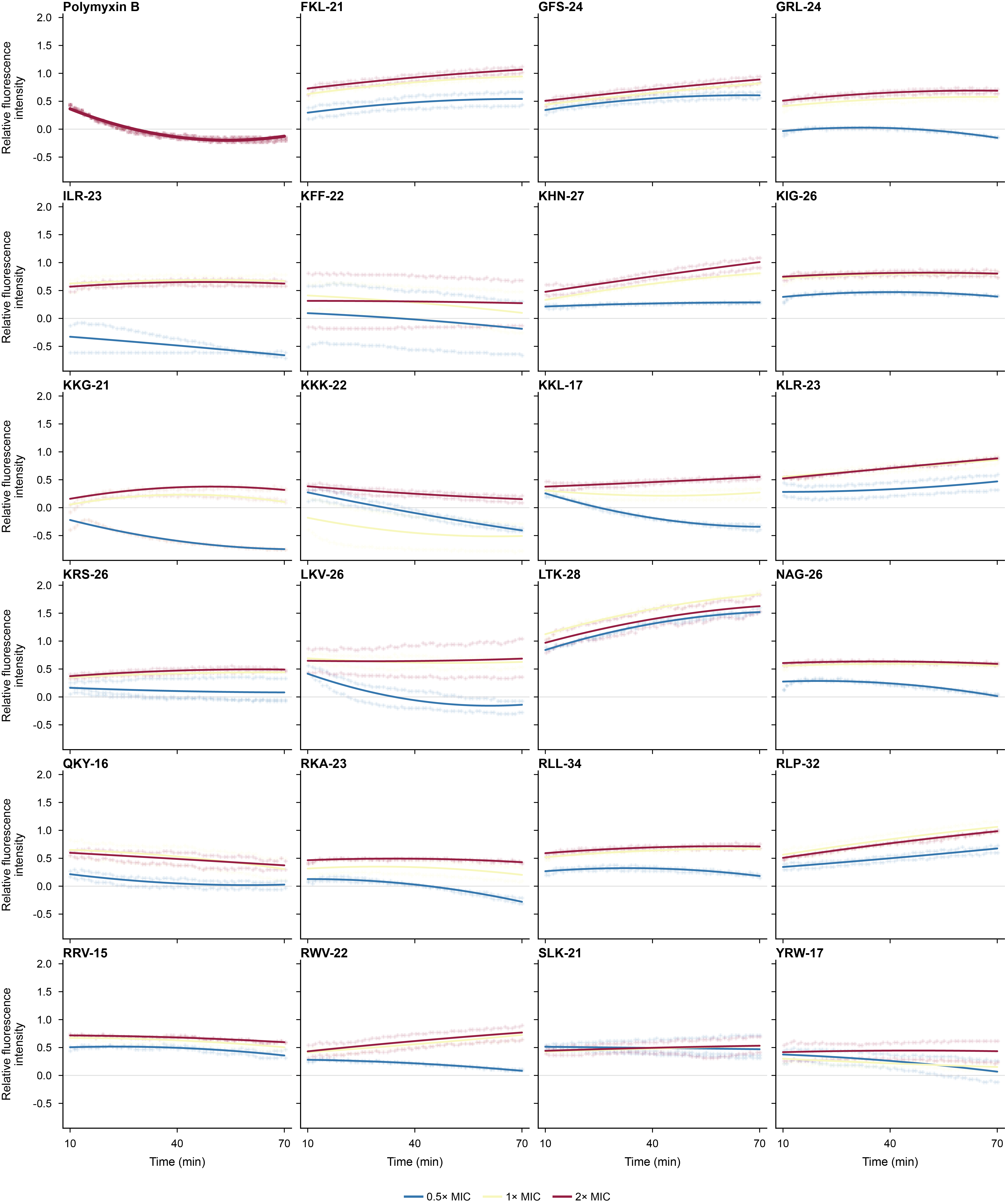

**Supplementary Figure 11 | Cytoplasmic-membrane depolarization of *Escherichia coli* measured using the membrane-potential-sensitive dye DiSC3(5).** Relative fluorescence was monitored for 60 min after treatment with the indicated peptides at 0.5× MIC, 1× MIC, and 2× MIC.

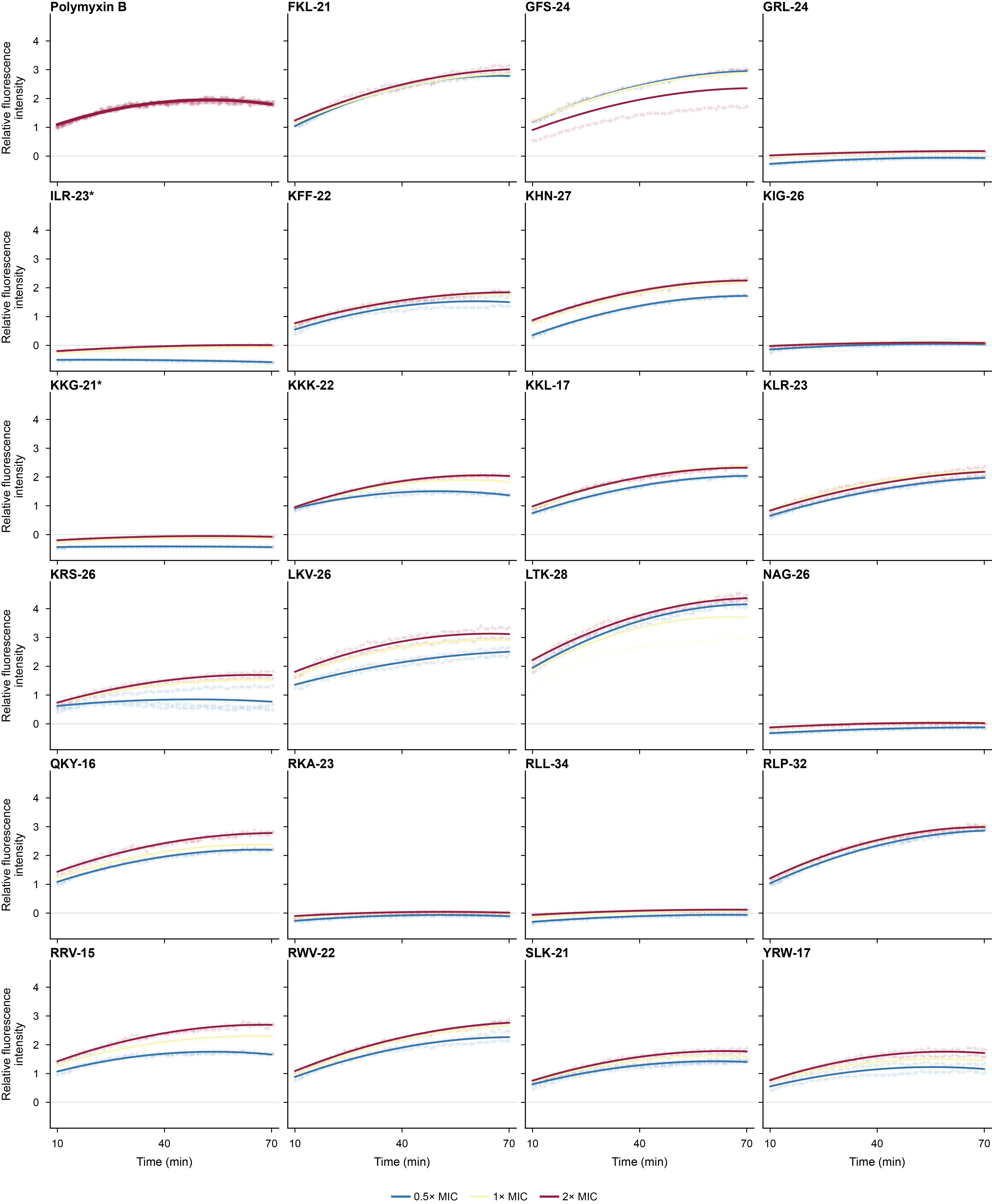

**Supplementary Figure 12 | Cytoplasmic-membrane depolarization of *Staphylococcus aureus* measured using the membrane-potential-sensitive dye DiSC3(5).** Relative fluorescence was monitored for 60 min after treatment with the indicated peptides at 0.5× MIC, 1× MIC, and 2× MIC.

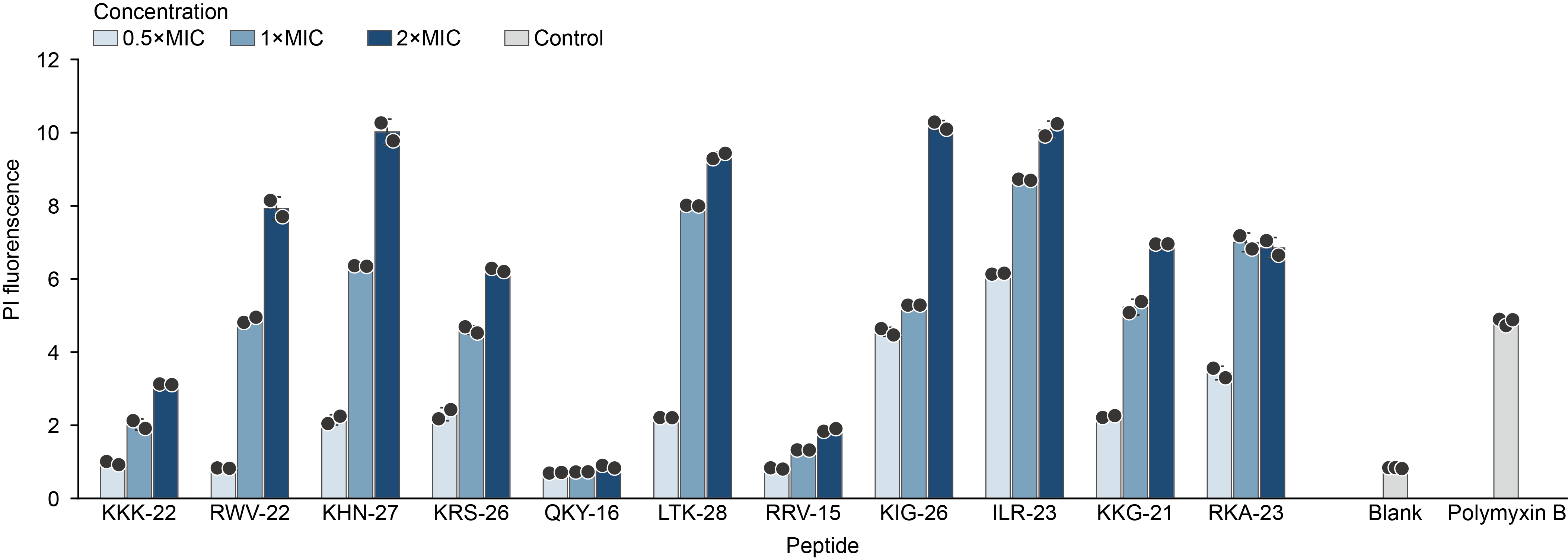

**Supplementary Figure 13 | PI uptake by *Escherichia coli*.** *Escherichia coli* was treated with the indicated peptides at 0.5×, 1× or 2× MIC. Blank and Polymyxin B were included as controls and are shown as grey bars. Bars and error bars show the mean and the sample standard deviation, respectively. Dots show individual measurements (n = 2 at each peptide concentration; n = 3 for Blank and Polymyxin B). An increase in PI fluorescence reports loss of cytoplasmic membrane integrity.

### 8. Cytotoxicity and haemolysis of the validated AMPs

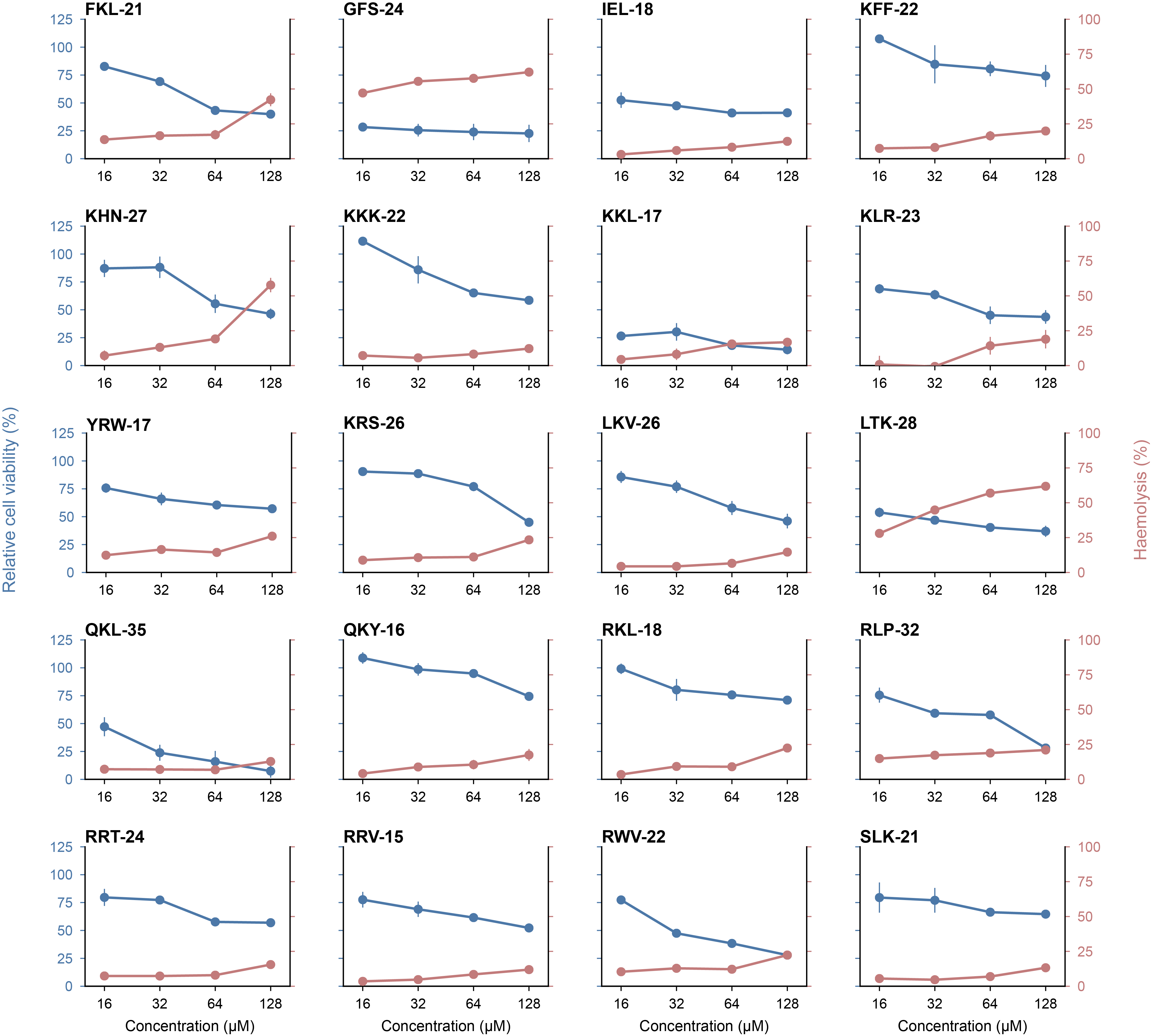

**Supplementary Figure 14** | **Dose-dependent cytotoxicity and hemolysis of the tested peptides.** Blue, relative cell viability (left axis); red-brown, hemolysis (right axis). Peptides were assayed at 16, 32, 64 and 128 µM. Points are the mean of two independent replicates; bars span the two replicate values. Viability is expressed relative to untreated cells (100%); hemolysis is expressed relative to complete lysis (100%).

### 9. Serum cytokine responses

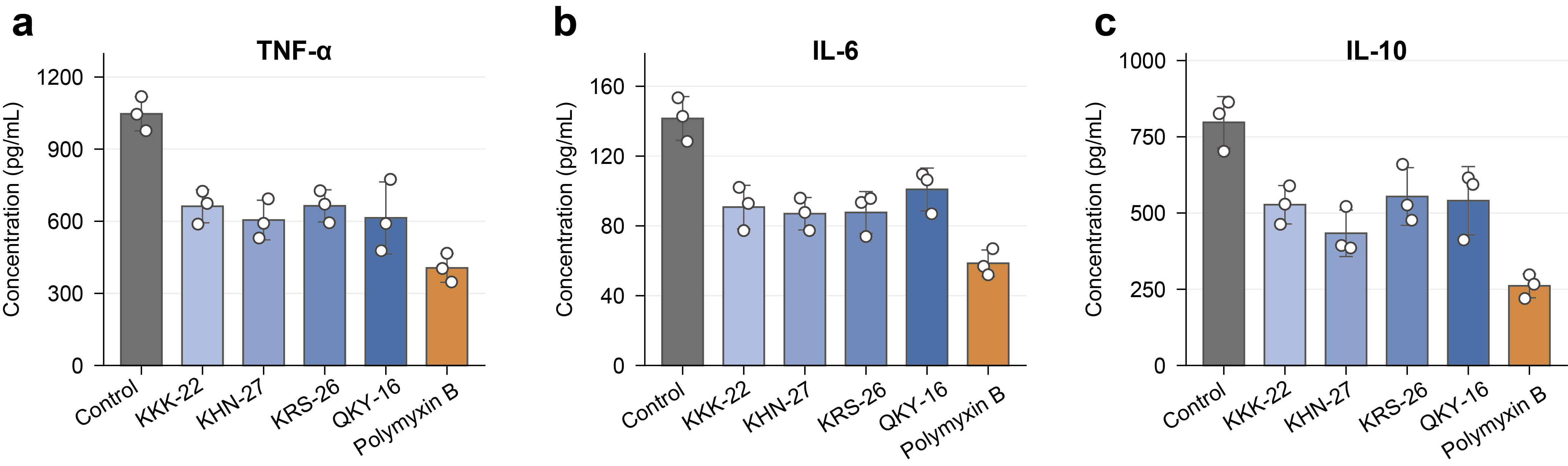

**Supplementary Figure 15** | **Serum cytokine levels following peptide or antibiotic treatment in a mouse muscle infection model.** Serum concentrations of TNF-α (**a**), IL-6 (**b**) and IL-10 (**c**) measured by ELISA in the control group and mice treated with KKK-22, KHN-27, KRS-26, QKY-16 or Polymyxin B. Open circles indicate individual measurements.
